# EntomonVR: a New Virtual Reality Game for Learning Insect Morphology

**DOI:** 10.1101/2023.02.01.526587

**Authors:** Mikaeel Pasandideh Saqalaksari, Ali Asghar Talebi, Thomas van de Kamp, Sajjad Reyhani Haghighi, Dominique Zimmermann, Adrian Richter

## Abstract

In recent years, the study of insect morphology has benefited greatly from the emergence of new digital imaging and analysis technologies such as X-ray micro-computed tomography (μ-CT), digital 3D reconstruction, and animation. Through interactive gaming and virtual reality, the external morphology of insects can be studied by a broad audience of both entomologists and non-specialists. EntomonVR is a serious game designed to investigate the external morphology of insects with adequate quality for the virtual reality platform. We discuss the advantages of virtual reality, introduce the EntomonVR new educational game, and conclude about future perspectives, validations, and cost-effectiveness. For game assessment, we have tested this game on 25 participants with an entomological background and improved the game based on their feedback. This study demonstrates the efficacy of virtual reality technology for an experimental learning environment in teaching the morphology of insects and the crucial needs for advancing an efficient and interactive educational program.

## Introduction

Insect morphology was a burgeoning discipline in the 20th century, with eminent works by insect researchers at many institutions (Matsuda, 1970; Snodgrass, 1939). From the beginning of the 21st century, entomologists have increasingly employed modern digital imaging, processing, and analysis techniques with the aim of generating detailed 3D reconstructions of specimens. The most prevalent 3D imaging technologies used by biologists are X-ray microcomputed tomography (Micro-CT) (Faulwetter et al., 2013; Hita Garcia et al., 2017; van de Kamp et al., 2018) and photogrammetry (Gontard et al., 2016; Gutiérrez-Heredia et al., 2015; Qian et al., 2015; Salle et al., 2014; Sosa et al., 2016; Ströbel et al., 2018).

The emergence of 3D visualisation tools has further contributed to the availability of virtual 3D specimens. Experiential learning is probably the most important method in the process of entomological education (Campo and Dangles, 2020). While it is based on physical specimens, hands-on experiments for students are generally rare. Virtual Reality (VR) is a cutting-edge technology that facilitates experiential learning (Piovesan et al., 2012) by allowing interaction with a simulated virtual world (Hamrol et al., 2013; Zilverschoon et al., 2017). A Virtual World is an artificial world made up of interactive graphical 3D models. This environment is accompanied by immersive visualisation techniques and tools for interaction and manipulation instantly in this artificial world (Bergmann et al., 2017; Jensen and Konradsen, 2018). Virtual reality systems come in many different types, for example, commercial and business systems available to the public, or more advanced systems that only some research centres and large companies have access to (Jensen and Konradsen, 2018).

Many educational programmes employ virtual reality technologies to overcome the limitations of 2D content in education and provide appropriate content for professional practice (Calvelo et al., 2020; Cao and Cerfolio, 2019; Cassidy et al., 2020; Ferrell et al., 2019). The interactive computer-generated experience can educate students by displaying things that are difficult or even impossible to observe, such as microscopic objects or the behaviour of organs in biological systems. This allows users to interact with an otherwise inaccessible environment in real-time. Additionally, virtual reality can be beneficial in education and learning for people with lower visual-spatial ability (Cui et al., 2017; Molina-Carmona et al., 2018).

In this study, we designed an interactive VR game in the field of entomology to provide an intuitive and attractive tool for students in the field of biology to enhance the learning experience. EntomonVR will be a useful tool for specialists and the non-specialist community. This allows experts to better investigate the external anatomy of insects and will be a visually striking and knowledge-based representation of insects in the virtual world for non-specialists. It also allows curators and communication programme coordinators in natural history museums to share informative 3D insects with students and the public.

## Method and Materials

### Hardware

The game framework uses the Oculus Rift CV1 headset to create a user environment in a 3D space for interaction within game activities. Technical specifications of the computer system for testing in the project were as follows: An Asus Rog G20BM mini-tower PC with a 770k Quad Core processor 3.50 GHz, Graphic Card Asus PH-GTX 1050 TI, and 16 GB RAM.

### Species 3D data availability

Details for each 3D data used on this platform are given in Table 1. The micro-CT images were reconstructed in FEI Amira 6.1 software (“Amira Software,” 2016). In Amira, the polygon of the models was generated by the Isosurface plugin and Extract surface tools. Afterwards, the extracted models were imported into MeshLab software (Cignoni et al., 2008) for post-processing. The post-processing steps followed the method of Hita Garcia et al., (2017) and subsequently, reassembling was done in Blender 2.82 software (“Blender3D software,” 2018). To optimise the models for the best performance and adequate details, 3D models with different ranges of polygons were developed (Figure 1).

**Table 1.**
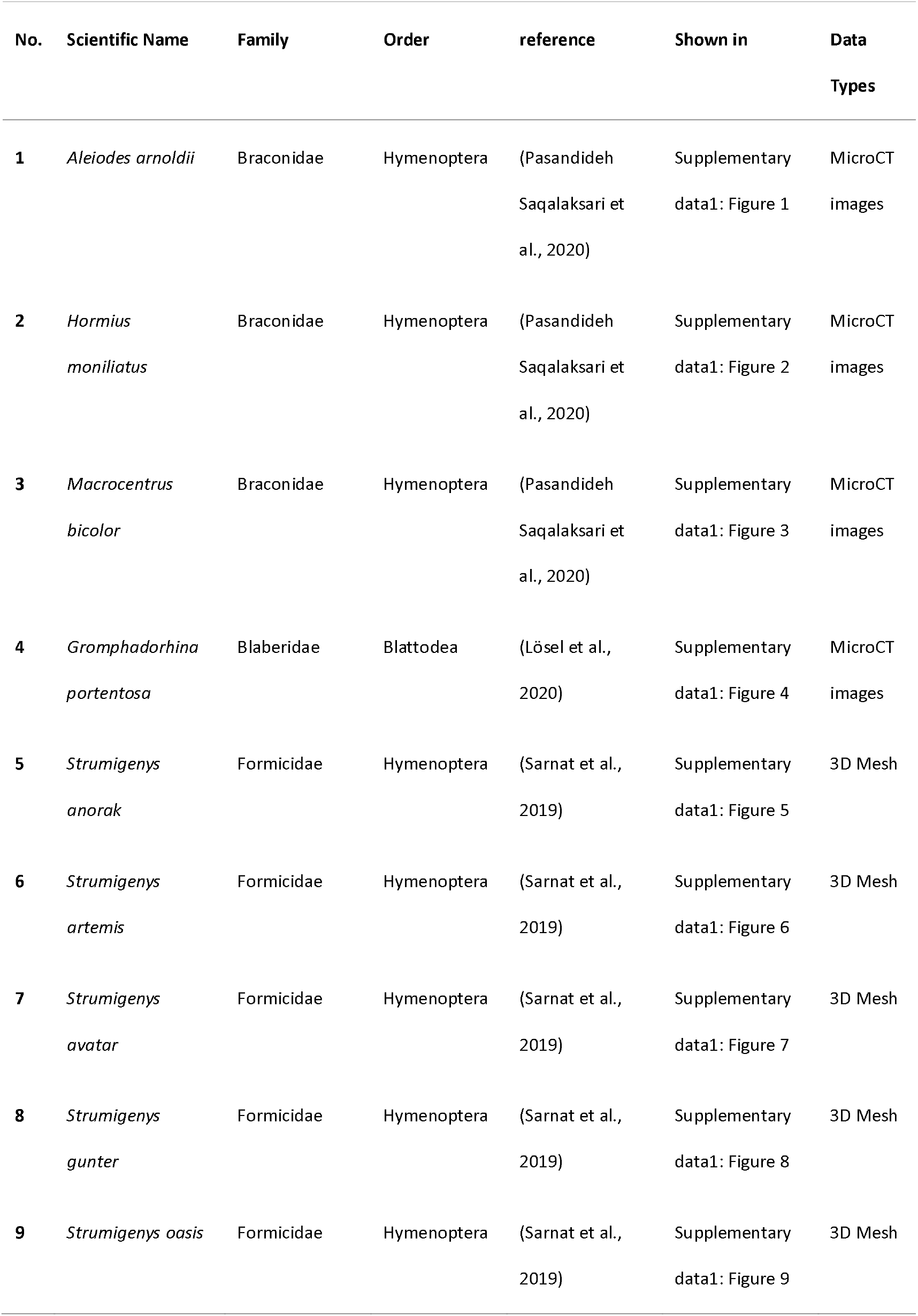
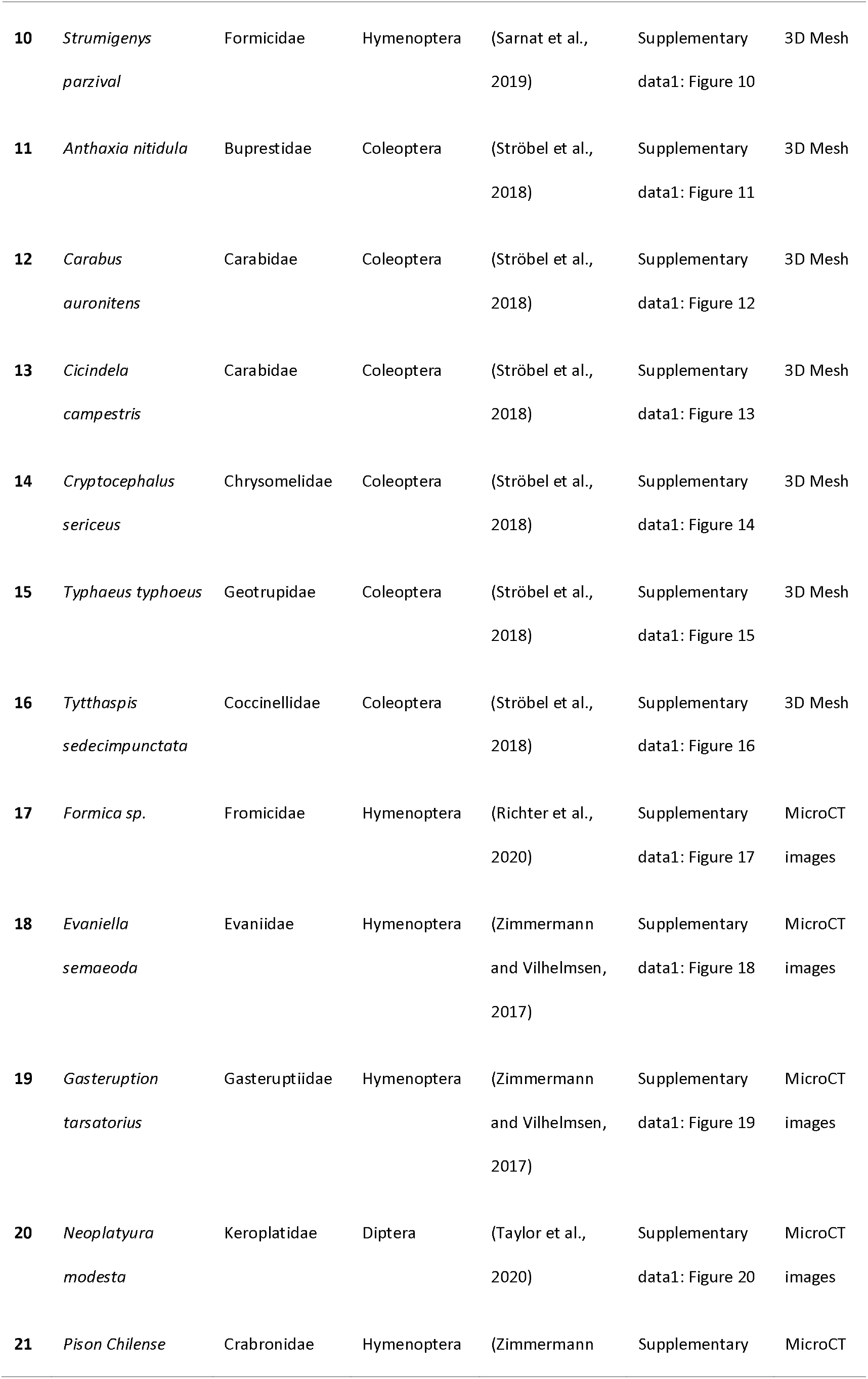
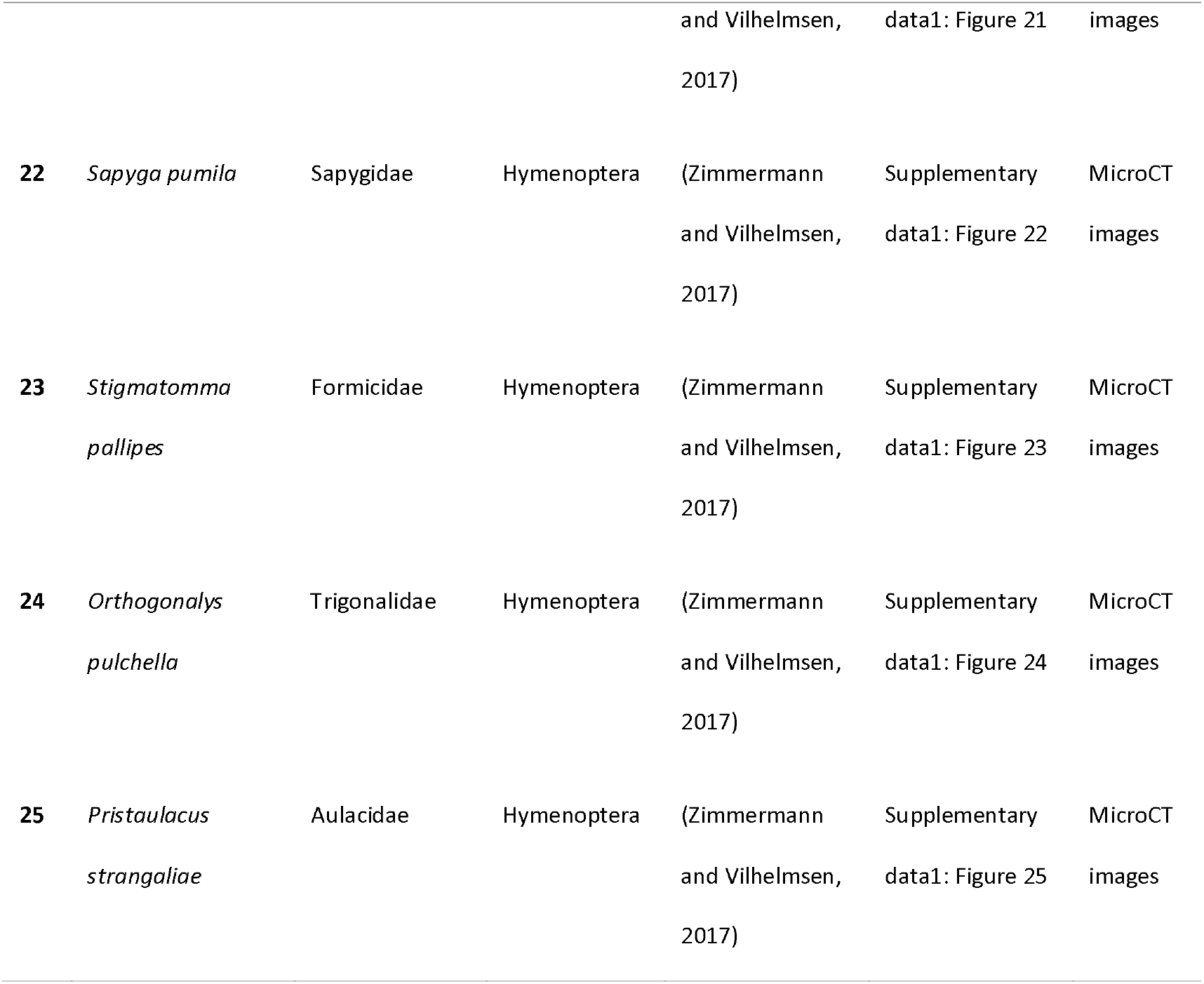
List of insect data’s used in the Basic version of EntomonVR

**Figure 1.**
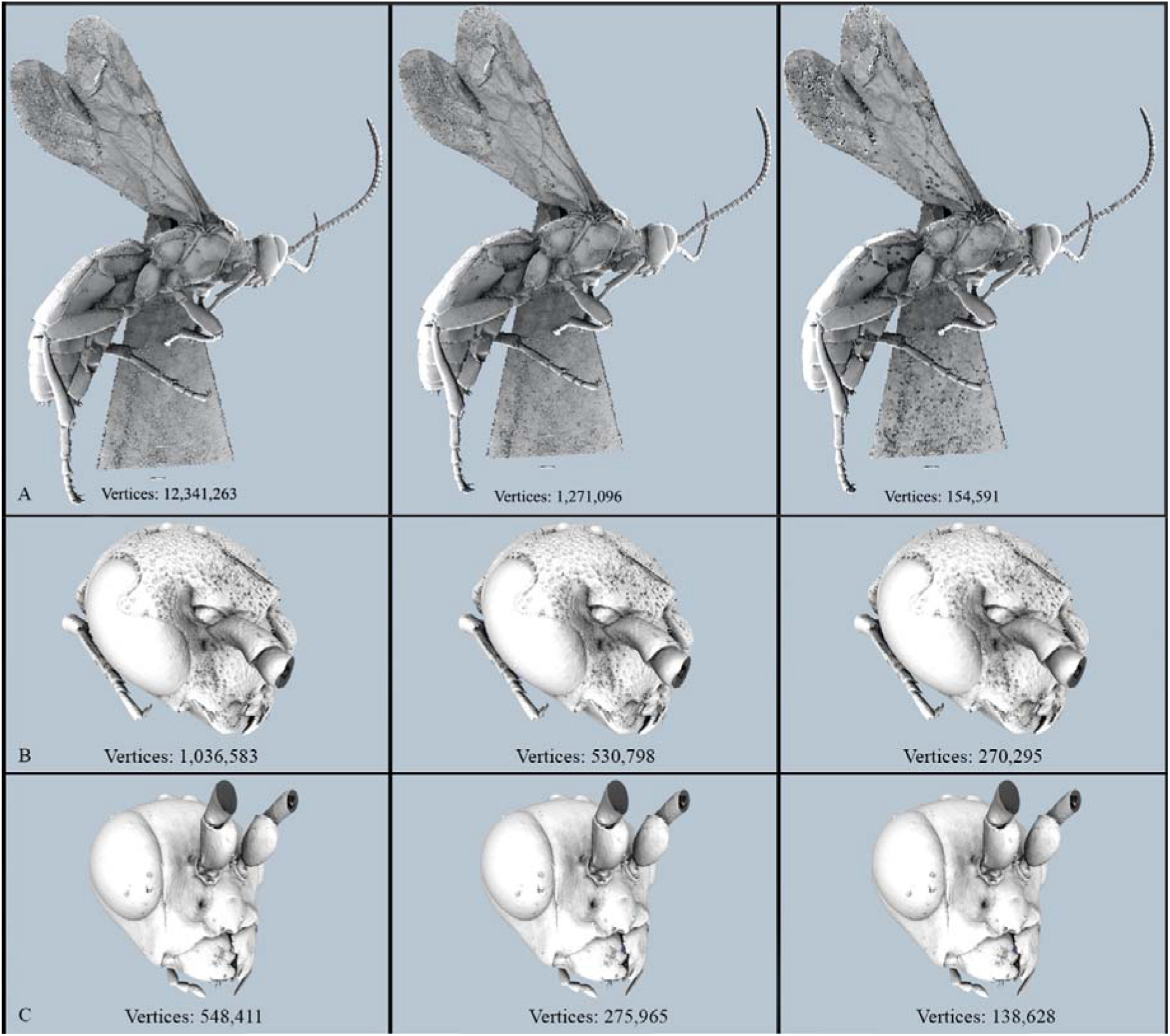
Surface rendered with different range of polygons for optimisation in virtual reality platform. A. *Aleiodes arnoldii* Tobias, 1976 (Hymenoptera: Braconidae); B. *Sapyga pumila* Cresson, 1880 (Hymenoptera: Sapygidae); C. *Pristaulacus strangaliae* Rohwer, 1917 (Hymenoptera: Aulacidae)

The 3D models reconstructed from CT volumes were painted based on their surface reference color with Vertex-Paint in Blender 2.82 (Figure 2) (Pasandideh Saqalaksari and Talebi, 2018).

**Figure 2.**
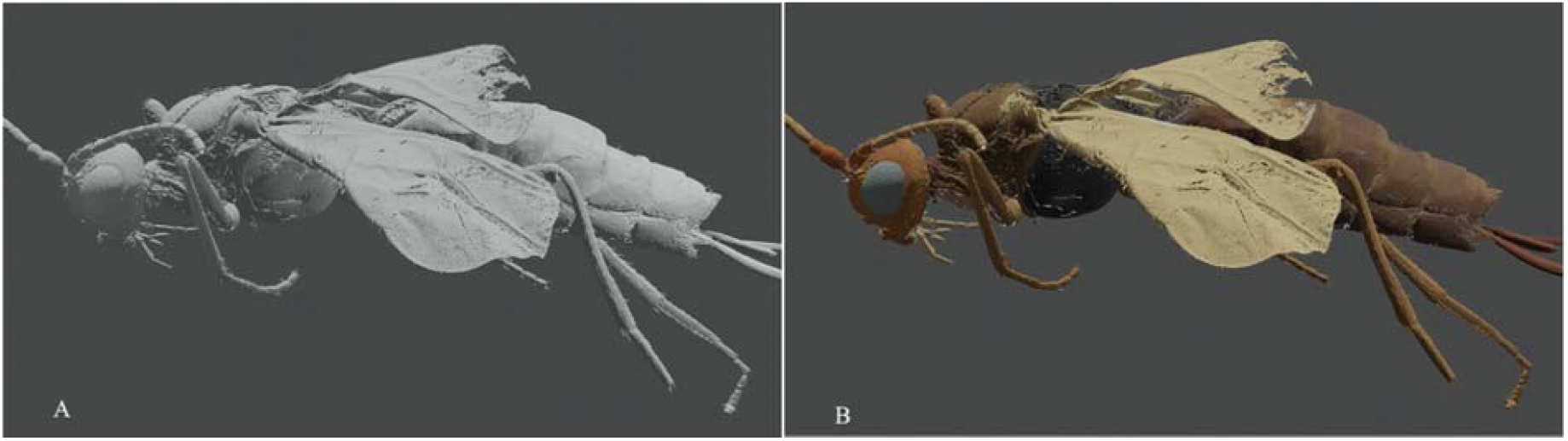
*Hormius Moniliatus* Nees, 1811 (Braconidae: Hormiinae) presented as an optimised 3D mesh model for virtual reality. A. Volumetric surfaces rendered model and B. Vertex painted colorised model from standard specimen images.

### Game development

Using the Unity3D 2018 game engine, the EntomonVR game was developed in the early prototype stage in which the player makes a journey through a designed level and interacts with different species. The digital 3D surface models were designed with Blender; the images were created with Adobe Photoshop CC 2018 (“Adobe Photoshop software,” 2018). Afterwards, the model has been exported to Unity3D (“Unity Software,” 2018). C# programming language was used to create the EntomonVR. The information such as 2D images and features description was gathered from different repositories and references (Beutel et al., 2016; Bolton, 1994; Yoder et al., 2010). An overview of the morphological characters of some insects examined in this study have been demonstrated in Figure 3, Figure 4, and Figure 5.

**Figure 3.**
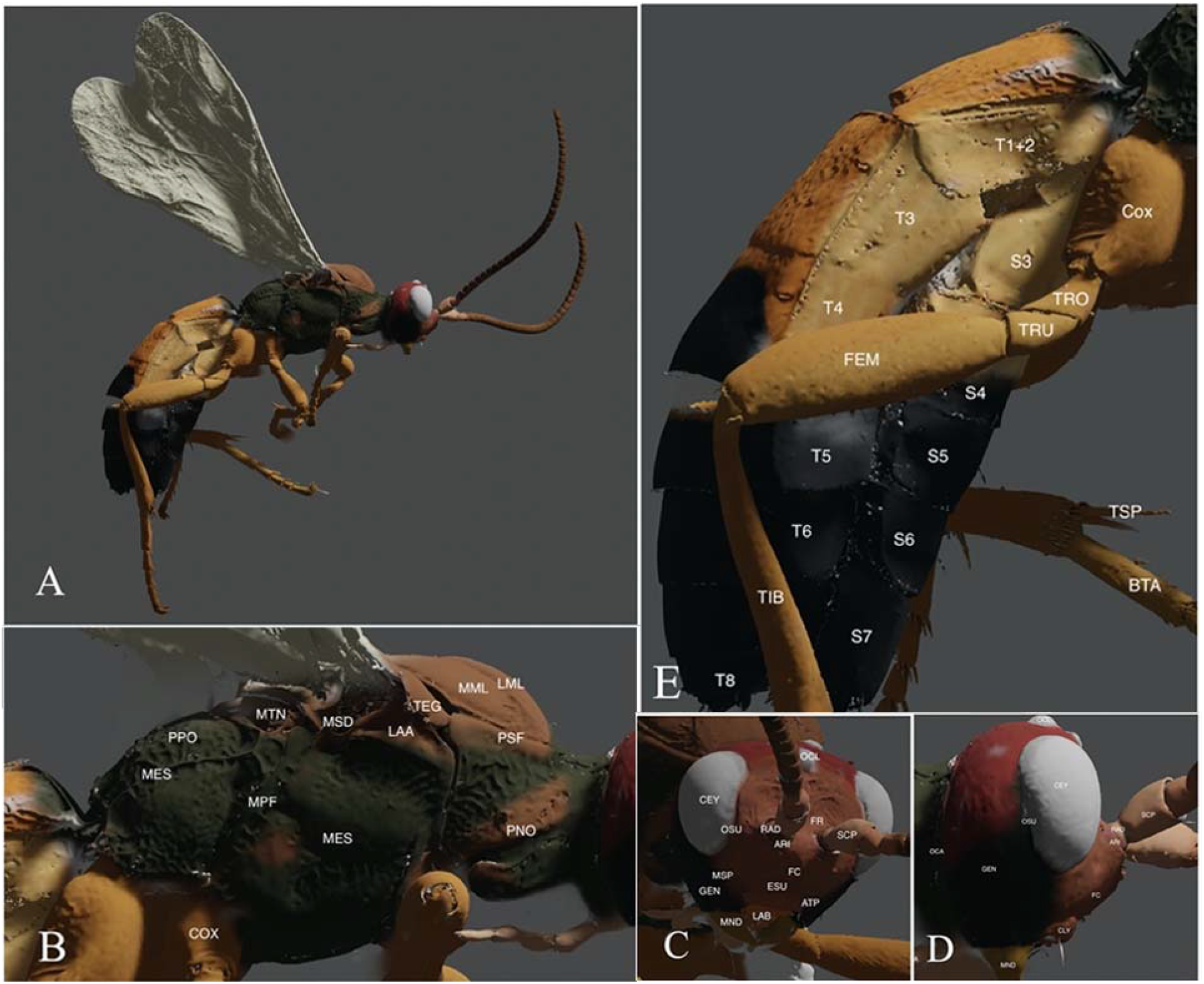
Details in 3D model of *Aleiodes arnoldii* Tobias, 1976 (Braconidae: Rogadinae); A. Lateral habitus, B. Thorax lateral view, C. Head frontal view, D. Head lateral view, E. Abdomen lateral view. Abbrevations: ARI, antennal rim; ATP, anterior tentorial pit; BTA, basitarsus; CEY, compound eye; Cox, coxa; CLY, clypeus; ESU, epistomal sulcus; FC, face; FEM, femur; FR, frons; GEN, gena; LAA, lateral axillar area; LAB, labium; LML, lateral mesoscutal lobes; MES, mesepisternum; MML, median mesoscutal lobe; MND, mandible; MPF, metanotal propodeal fissure; MSO, mesonotum; MSP, malar space; MTN, metanotum; OCA, occipital carina; OCL, ocellus; OSU, ocellar suture; PNO, pronotum; PPO, propodeum; PSF, parascutal flange; RAD, radicle; S3-S7, abdominal sterna 3-7; SCP, scape; T1+2, abdominal tergum 1+2; T3-T8, metosomal tergum 3-8; TEG, tegula; TIB, tibia; TRO, trochanter; TRU, protrochantellus; TSP, tibial spore.

**Figure 4.**
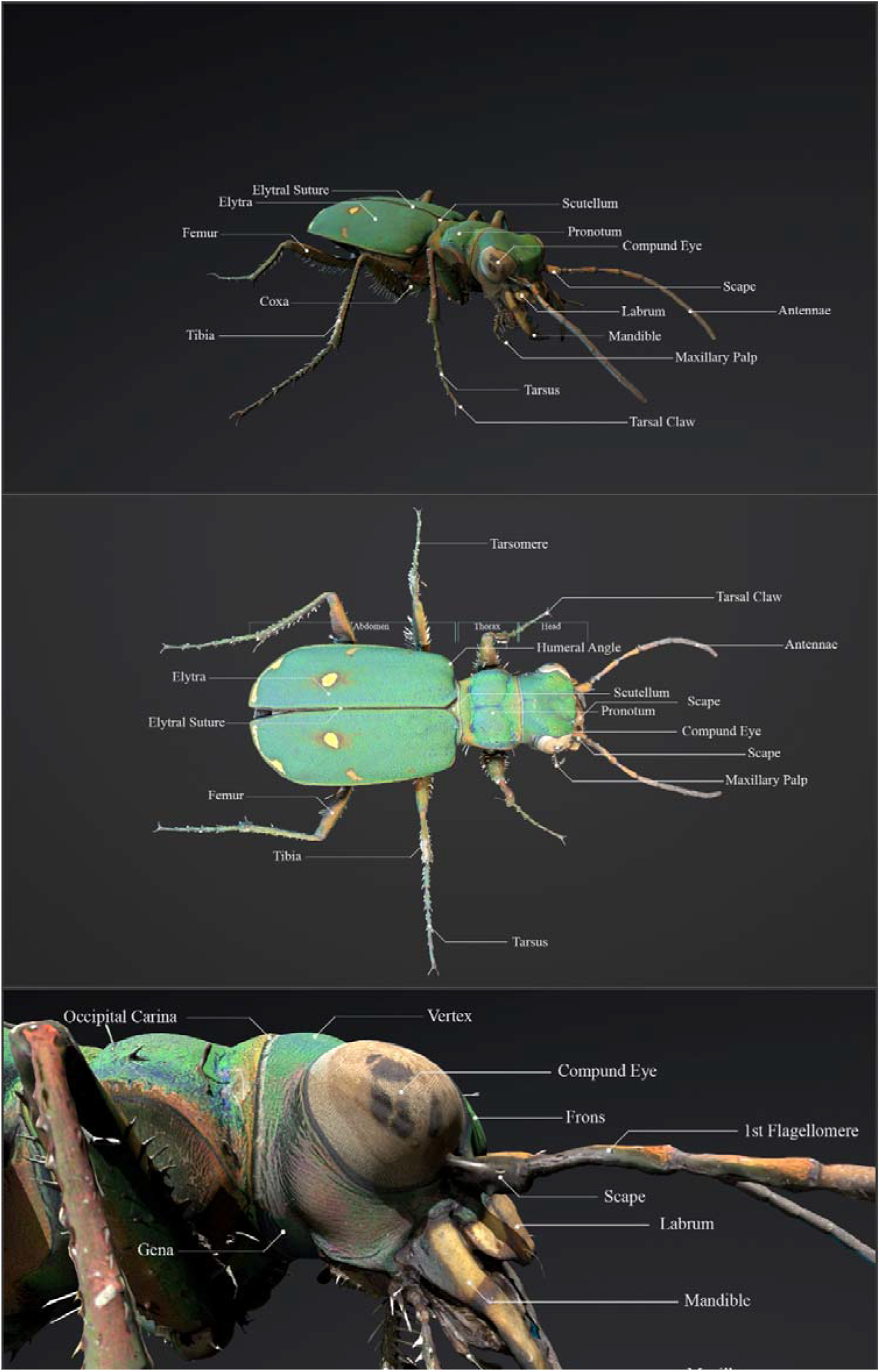
Details in 3D model of *Cicindela campestris* Linnaeus, 1758 (Coleoptera: Cicindelidae); A. Whole body frontal view, B. Whole body dorsal view, C. Head lateral view.

**Figure 5.**
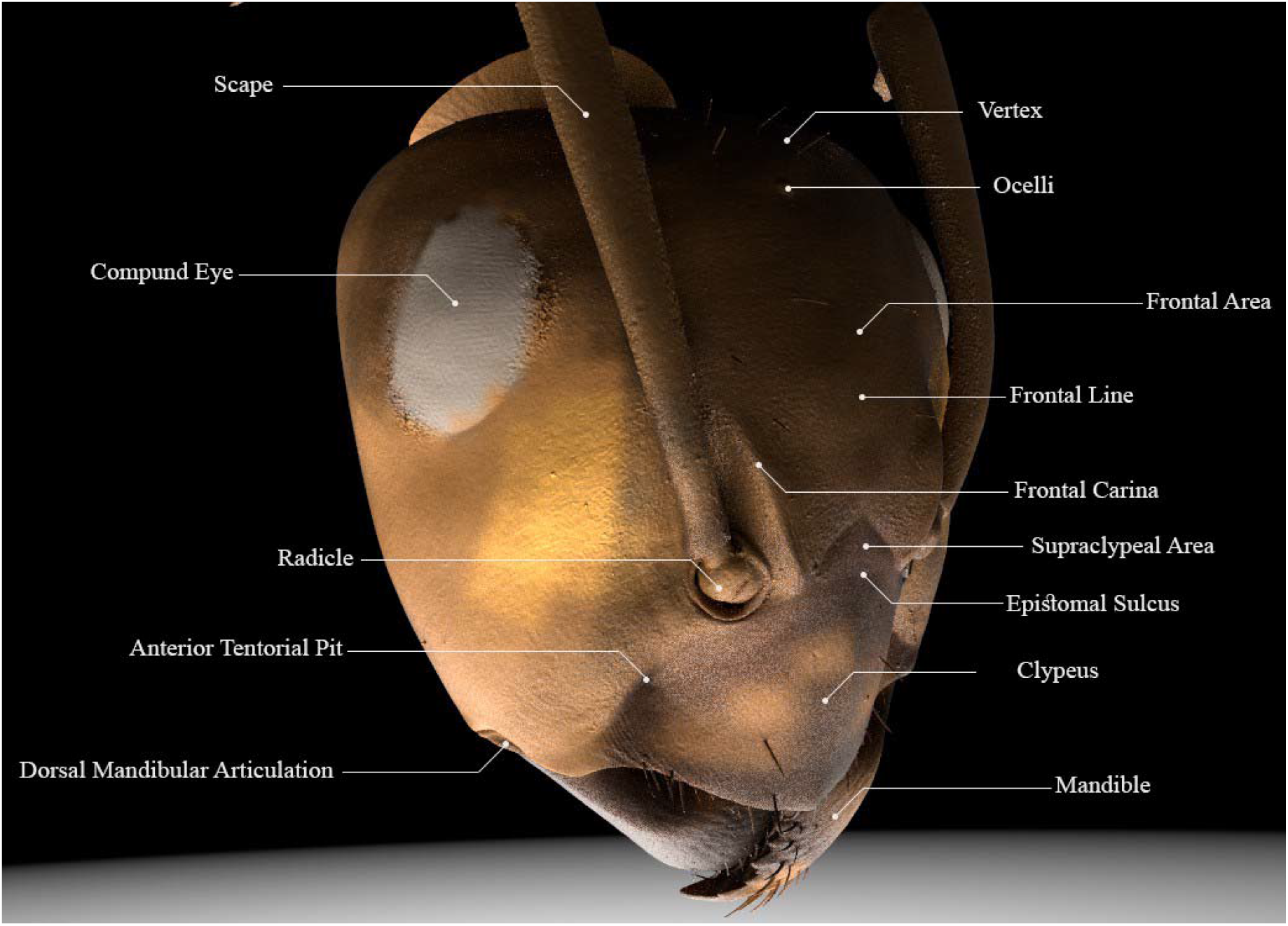
Details in 3D model of the head *Formica sp*. (Hymenoptera: Formicidae).

The current framework was developed through a series of interviews with students and scientific staff. All interested groups understood the concept and issued specific requirements like intangible characters and simple UI to facilitate the operations. Students with an entomological background were selected for preliminary tests. These volunteers provided their feedback about the game to the technical team for the necessary development.

### Prototype assessment

To evaluate and improve the EntomonVR game, 25 participants (10 males, 15 females with age range of 21 to 30 years old) with entomological knowledge from Tarbiat Modares University, University of Tehran, Shahid Beheshti University and Guilan University were asked to play the game and to complete the post-study questionnaire about their experiences.

Three of the 25 participants had previous VR gaming experience. The participants completed an integrated questionnaire about their perceptions of the technology, object reality, and scientific information as feedback for possible improvements of the game-based framework. The questionnaire consisted of 30 questions based on a 5-star rating (Supplementary data 2).

## Results

The EntomonVR is available for free on Mendeley Data (doi:10.17632/jfx5zsypxf.1) and is compatible with all VR headsets. EntomonVR allows users to experience 3D models of insects. The current version features 25 species from four different orders of insects (Table 1). The virtual entomology lab was implemented, where models of all 25 species were placed and users select which species to look at using the specimen selector button. The models can be grabbed using the controller, scaled and moved around individually (Figure 6A). For getting a good overview of body parts, info-boxes have been set for each body segment to show information of each part in the virtual tablet (Figure 6B). By scaling the models, the smaller characters appear with info-rings. Users can interact with these and view related information in the tablet (Figure 6C). For navigation in the virtual lab, a new teleport navigation scheme is selected with hand controllers, the VR-controller joystick (Figure 7) and during this process, a green arc is displayed that enables movement to the desired point in the environment (Figure 6D). A video demonstrating the features and user guide of EntomonVR is available from the online version of this article (Supplementary data 3) and on YouTube (https://youtu.be/sP2gGsmjxqo).

**Figure 6.**
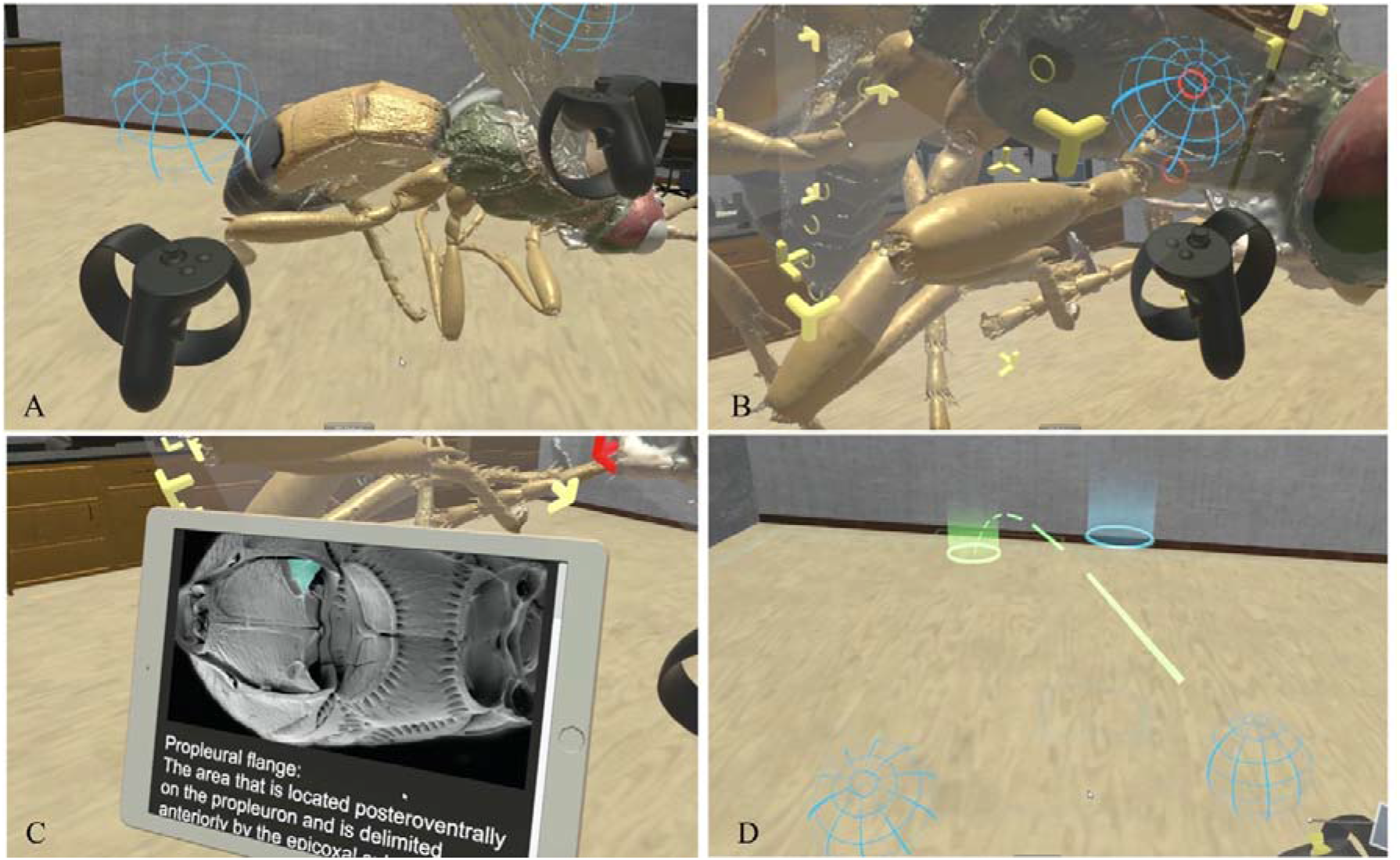
Screenshots are demonstrating the EntomonVR app. The EntomonVR is available for free on Mendeley Data (doi:10.17632/jfx5zsypxf.1) and is compatible with all VR headsets. A video demonstrating the features and user guide of EntomonVR is available in the online version of this article (Supplementary data 3) and on YouTube (https://youtu.be/sP2gGsmjxqo). A. 3D view of the cyber specimen. B. InfoSet and InfoRing are demonstrating the morphological characters C. Virtual tablet displaying characters images and related information. (D). The teleporter in the virtual lab.

**Figure 7.**
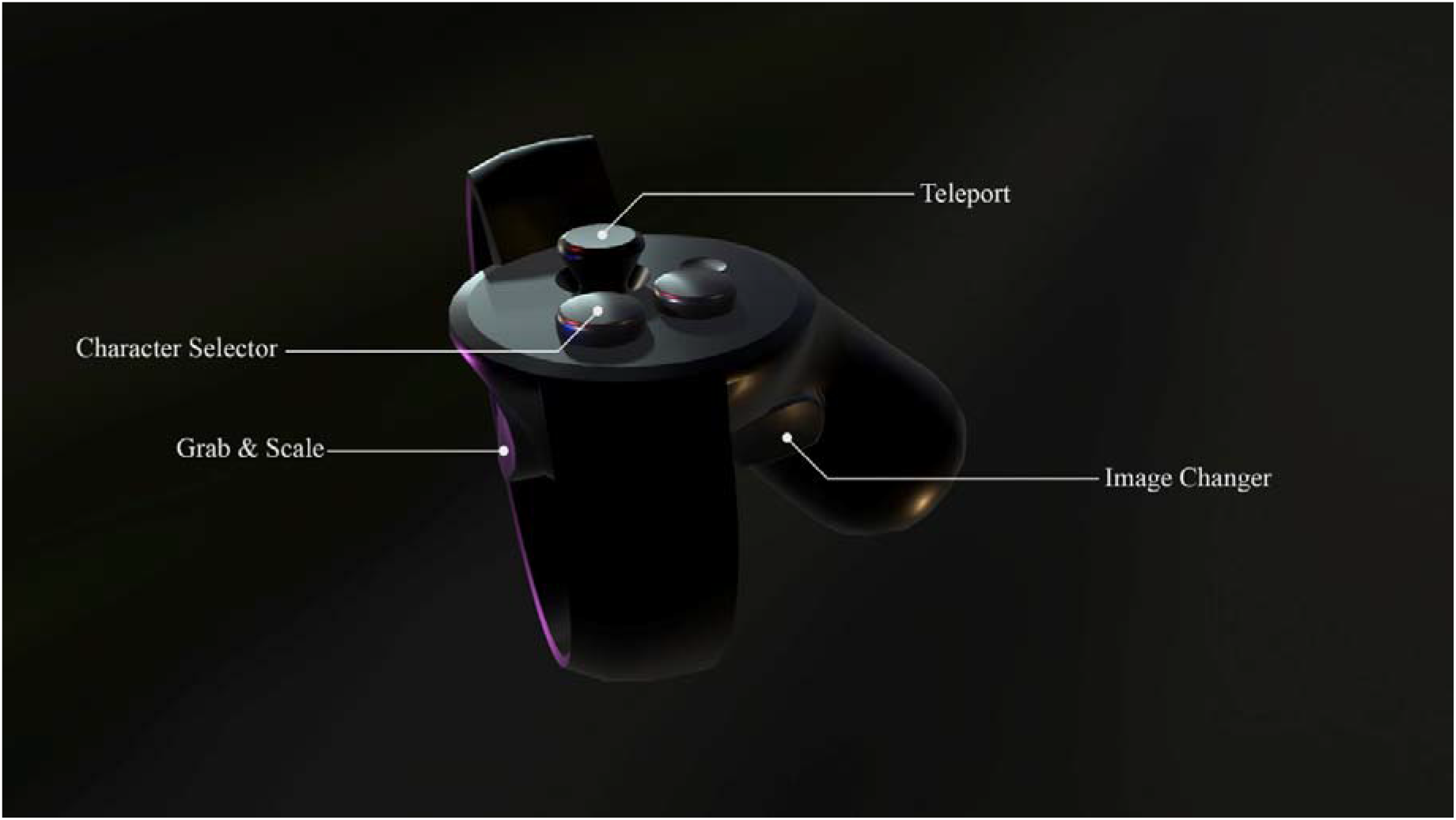
Illustration of the VR controller buttons with different functions (Oculus Touch Controller)

The majority of participants had no experience in VR gaming which led to the development of a tutorial guide. Three participants had previous VR gaming experience in the VR clubs and the 23 other participants encountered problems with the game and performed with technician assistance.

Overall, all the answers were within the high score that 33.75% of the answers were five stars, 30.56% of the answers four stars, 14.72% of the answers three stars, 2.78% of the answers two stars and 1.11% of the answers were one star. This means that volunteers were attracted to the framework, and they provided our research team with some suggestions for colouring the insect models, adding more insects from different orders to the game, and more interactable insect elements (such as internal organs and parasitic organisms). Moreover, adding instructions in the gameplay and tutorial level to familiarise with the VR technology were suggested and some participants suggested the Q&A step in the game. The majority of the participants were happy with the game-based framework and believed it perfectly showed the insect body parts. Two participants wanted to make storyboards for upcoming developments and an entire game scenario.

The academic staff stated that students would be excited about the use of virtual reality within the laboratory setting. 50% of the participants stated the option to interact with different parts of each insect increased the quality of learning noticeably. They also provided ideas for further development of specific game challenges.

## Discussion

With the increasing expansion of virtual reality headsets and VR learning games, EntomonVR provides a new option for virtual reality learning to immerse users in touch and feel the insect’s body and get to know more about these organisms. Interaction is an integral part of game-based learning for students with varying interactable objects (Plass et al., 2015). Along with the significant improvement of game engines such as Unreal Engine and Unity3D, more appropriate tools are available for developers to create convenient and stable virtual environments.

Traditional methods used in entomological education and learning are associated with some potential issues as they can provide a limited understanding obtained from oral lectures and restricted anatomical dissection (Leonard and Penick, 2000). 3D reconstruction and modelling have enabled us to visualise structures from different viewpoints, change the size, and annotate explorative details and thus have opened up a new view for education.(Brenton et al., 2007). In this study, these issues are overcome by innovative imaging technologies involving micro-computed tomography and other three-dimensional visualisation methods. By changing to the utilisation of 3D technologies in learning, it is also becoming necessary to align these technologies to accepted assessment methods (Bogomolova et al., 2021). While the shift in education has transposed from a two-dimensional nature to three-dimensional methods, there is a need to change assessment methods in a parallel direction and proceed towards using three-dimensional and emerging technologies.

Virtual reality can deliver real-world experiences into any place like schools, homes or social centres (Dorward et al., 2017). The results show that EntomonVR can be used by entomology students to recognise and understand the structures and characters, enabling them to obtain knowledge and better understand insect morphology by interacting within a virtual world. Among other advantages, this immersive technology provides the possibility to modify sizes in virtual space, whereas this would be impossible to accomplish in reality (Johnston et al., 2018).

The decrease in the number of vertices and faces in a 3D model is based on the reduction level that is the reduced percentage of vertices (Low and Tan, 1997). The purpose of this work is to prepare 3D scanned model for virtual reality and game engines. The proposed method significantly reduces the size of the model by maintaining the topology with high accuracy of faces. The reduction in the number of polygons in the models with sculpted surfaces is evident in high levels and causes the loss of some delicate structures such as setae and ommatidia. Nevertheless, in the models with smooth surfaces, this reduction can be achieved up to high levels with minimal amount of loss of details.

Many methods for the 3D digitisation of insect specimens have been developed, for example based on multiple photography (e.g. Ströbel et al., 2018) and computed tomography (Greco et al., 2008). Both approaches have some advantages and disadvantages. 3D reconstruction using photogrammetry obtains both shapes and textures simultaneously, but it is challenging to reconstruct concave structures (Salle et al., 2014). Computed tomography provides full anatomy accurately even if a specimen has concave structures (Greco et al., 2008). However, X-ray imaging cannot process surface colours, and surface texturing is a difficult action. Adding colour to 3D models can improve the engagement of the non-scientific audience and even researchers. Ijiri et al., (2018) presented a technique for digitising insects by combining Micro-CT scans and photogrammetry to create 3D textured models. Although 360-degree image-based techniques are suggested, using these methods for hymenopteran winged specimens reveals obvious drawbacks described by Ströbel et al., (2018). Therefore, in this study, the vertex-paint method was used and the participants found it satisfactory.

Despite difficulties with controllers, game-based learning is supported by initial users and a list of comments for improvement of insect realistic elements is suggested (Supplementary data 02). Special interactions need new types of user interfaces. In this study, the specimen selector button was implemented to show the 3D models in the virtual lab and virtual tablet to browse the information of each character. We recommend using real-world interaction plans such as VR gloves and haptics in the VR environment to make the user’s learning process more comfortable.

## Conclusion

Virtual reality (VR) offers a range of new opportunities, particularly in the field of studying 3D cyber-specimens. The use of VR models, such as those of braconid wasps, can greatly enhance understanding of insect anatomy through precise and visually engaging representations. Additionally, the VR software can be utilised for educational and training purposes.

This study illustrates the significant impact that advancements in computer graphics technology are having, and will continue to have, on insect science. There are plans to further develop EntomonVR into a virtual insect laboratory, featuring a wider range of species and disciplines. VR headsets are powerful tools for visualizing 3D objects, providing high-quality and engaging scientific visualization.

## Author Contributions

M.P.S. designed the study, drafted the manuscript, created volume rendering and surface models, and developed the VR app. A.T. supervised the project. T.v.d.K. helped to supervise the project, performed microCT, and worked on the manuscript. S.R.H. contributed to the interpretation of the results and provided critical feedback. A.R. provided ant heads microCT data and related morphological information. D.Z. provided microCT data of some hymenopteran species and worked on the manuscript. A.L. provided critical review, contributed to the interpretation of the results and worked on the manuscript. All authors discussed the results and commented on the manuscript.

## References

Adobe Photoshop software, 2018.

Amira Software, 2016.

Bergmann, T., Balzer, M., Hopp, T., Van De Kamp, T., Kopmann, A., Jerome, N. T. & Zapf, M. (2017). Inspiration from VR gaming technology: Deep immersion and realistic interaction for scientific visualization. [Paper presentation] VISIGRAPP 2017 - Proceedings of the 12th International Joint Conference on Computer Vision, Imaging and Computer Graphics Theory and Applications. (e.g. https://doi.org/10.5220/0006262903300334”)

Beutel R. G., & Leschen, A. B. (2016). Handbook of zoology: Arthropoda: Insecta. Coleoptera, Beetles, vol 1. (2nd ed.). De Gruyter Press

Blender3D software, 2018.

Bogomolova, K., Sam, A. H., Misky, A. T., Gupte, C. M., Strutton, P. H., Hurkxkens, T. J., & Hierck, B. P. (2021). Development of a virtual three dimensional assessment scenario for anatomical education. Anatomical sciences education, 14 (3), 385–393. (e.g. “https://doi.org/10.1002/ase.2055”)

Bolton, B. (1994). Identification guide to the ant genera of the world. Harvard University Press.

Brenton, H., Hernandez, J., Bello, F., Strutton, P., Purkayastha, S., Firth, T., & Darzi, A. (2007). Using multimedia and Web3D to enhance anatomy teaching. Computers & Education, 49(1), 32–53. (e.g. “https://doi.org/10.1016/j.compedu.2005.06.005”)

Calvelo, M., Piñeiro, Á. & Garcia-Fandino, R. (2020). An immersive journey to the molecular structure of SARS-CoV-2: Virtual reality in COVID-19. Computational and structural biotechnology journal, 18, 2621–2628. (e.g. “https://doi.org/10.1016/j.csbj.2020.09.018”)

Campo, P., & Dangles, O. (2020). An overview of games for entomological literacy in support of sustainable development. Current opinion in insect science, 40, 104–110. (e.g. “https://doi.org/10.1016/j.cois.2020.05.018”)

Cao, C., & Cerfolio, R. J. (2019). Virtual or augmented reality to enhance surgical education and surgical planning. Thoracic surgery clinics, 29(3), 329–337. (e.g. “https://doi.org/10.1016/j.thorsurg.2019.03.010”)

Cassidy, K. C., Šefčík, J., Raghav, Y., Chang, A., & Durrant, J. D. (2020). ProteinVR: Web-based molecular visualization in virtual reality. PLoS computational biology, 16(3), e1007747. (e.g. “https://doi.org/10.1371/journal.pcbi.1007747”)

Cignoni, P., Callieri, M., Corsini, M., Dellepiane, M., Ganovelli, F., & Ranzuglia, G. (2008). MeshLab: An open-source mesh processing tool. [Paper presentation] 6th Eurographics Italian Chapter Conference 2008 - Proceedings.

Cui, D., Wilson, T. D., Rockhold, R. W., Lehman, M. N., & Lynch, J. C. (2017). Evaluation of the effectiveness of 3D vascular stereoscopic models in anatomy instruction for first year medical students. Anatomical sciences education, 10(1), 34–45. (e.g. “https://doi.org/10.1002/ase.1626”)

Dorward, L. J., Mittermeier, J. C., Sandbrook, C., & Spooner, F. (2017). Pokémon Go: Benefits, costs, and lessons for the conservation movement. Conservation Letters, 10(1), 160–165. (e.g. “https://doi.org/10.1111/conl.12326”)

Faulwetter, S., Vasileiadou, A., Kouratoras, M., Dailianis, T., & Arvanitidis, C. (2013). Micro-computed tomography: Introducing new dimensions to taxonomy. ZooKeys, (263), 1. (e.g. “https://doi.org/10.3897/zookeys.263.4261”)

Ferrell, J. B., Campbell, J. P., McCarthy, D. R., McKay, K. T., Hensinger, M., Srinivasan, R., Zhao, X., & Schneebeli, S. T. (2019). Chemical exploration with virtual reality in organic teaching laboratories. Journal of Chemical Education, 96(9), 1961–1966. (e.g. “https://doi.org/10.1021/acs.jchemed.9b00036”)

Gontard, L. C., Schierholz, R., Yu, S., Cintas, J. & Dunin-Borkowski, R. E. (2016). Photogrammetry of the three-dimensional shape and texture of a nanoscale particle using scanning electron microscopy and free software. Ultramicroscopy, 169, 80–88. (e.g. “https://doi.org/10.1016/j.ultramic.2016.07.006”)

Greco, M., Jones, A., Spooner-Hart, R., & Holford, P. (2008). X-ray computerised microtomography (MicroCT): a new technique for assessing external and internal morphology of bees. Journal of Apicultural Research, 47(4), 286–291.

Gutiérrez□ Heredia, L., D’Helft, C. & Reynaud, E. G. (2015). Simple methods for interactive 3D modeling, measurements, and digital databases of coral skeletons. Limnology and Oceanography: Methods, 13(4), 178–193. (e.g. “https://doi.org/10.1002/lom3.10017”)

Hamrol, A., Górski, F., Grajewski, D. & Zawadzki, P. (2013). Virtual 3D atlas of a human body - Development of an educational medical software application [paper presentation] Procedia Computer Science. (e.g. “https://doi.org/10.1016/j.procs.2013.11.036”)

Garcia, F. H., Fischer, G., Liu, C., Audisio, T. L., & Economo, E. P. (2017). Next-generation morphological character discovery and evaluation: an X-ray micro-CT enhanced revision of the ant genus *Zasphinctus* Wheeler (Hymenoptera, Formicidae, Dorylinae) in the Afrotropics. ZooKeys, (693), 33. (e.g. “https://doi.org/10.3897/zookeys.693.13012”)

Ijiri, T., Todo, H., Hirabayashi, A., Kohiyama, K., & Dobashi, Y. (2018). Digitization of natural objects with micro CT and photographs. Plos one, 13(4), e0195852. (e.g. “https://doi.org/10.1371/journal.pone.0195852”)

Jensen, L., & Konradsen, F. (2018). A review of the use of virtual reality head-mounted displays in education and training. Education and Information Technologies, 23(4), 1515–1529.

Johnston, A. P., Rae, J., Ariotti, N., Bailey, B., Lilja, A., Webb, R., McGhee, J. & Parton, R. G. (2018). Journey to the centre of the cell: Virtual reality immersion into scientific data. Traffic, 19(2), 105–110. (e.g. “https://doi.org/10.1111/tra.12538”)

Leonard, W. H., & Penick, J. E. (2000). The Limits of Learning. The American Biology Teacher, 62(5), 359–361. (e.g. “https://doi.org/10.2307/4450919”)

Lösel, P. D., van de Kamp, T., Jayme, A., Ershov, A., Faragó, T., Pichler, O., … & Heuveline, V. (2020). Introducing Biomedisa as an open-source online platform for biomedical image segmentation. Nature communications, 11(1), 1–14. (e.g. “https://doi.org/10.1038/s41467-020-19303-w”)

Low, K. L., & Tan, T. S. (1997). Model simplification using vertex-clustering. [Paper presentation] Proceedings of the 1997 symposium on Interactive 3D graphics (pp. 75–ff). (e.g. “https://doi.org/10.1145/253284.253310”)

Matsuda, R. (1970). Morphology and evolution of the insect thorax. The Memoirs of the Entomological Society of Canada, 102(S76), 5–431. (e.g. “https://doi.org/10.4039/entm10276fv”)

Molina-Carmona, R., Pertegal-Felices, M. L., Jimeno-Morenilla, A., & Mora-Mora, H. (2018). Virtual reality learning activities for multimedia students to enhance spatial ability. Sustainability, 10(4), 1074. (e.g. “https://doi.org/10.3390/su10041074”)

Pasandideh Saqalaksari, M., & Talebi, A. A. (2018). Iranian Braconidae. http://cyberbraconid.myspecies.info/ [Accessed on 2022/12/26].

Pasandideh Saqalaksari, M., Talebi, A. A., & van de Kamp, T. (2020). MicroCT 3D reconstruction of three described braconid species (Hymenoptera: Braconidae). Journal of Insect Biodiversity and Systematics, 6(4), 331–342.

Piovesan, S. D., Passerino, L. M., & Pereira, A. S. (2012). Virtual Reality as a Tool in the Education. [Paper presentation] International Association for Development of the Information Society (IADIS) International Conference on Cognition and Exploratory Learning in Digital Age (CELDA) (Madrid, Spain, Oct 19-21, 2012).

Plass, J. L., Homer, B. D. & Kinzer, C. K. (2015). Foundations of game-based learning. Educational psychologist, 50(4,: 258–283. (e.g. “https://doi.org/10.1080/00461520.2015.1122533”)

Qian, J., Lei, M., Dan, D., Yao, B., Zhou, X., Yang, Y., Yan, S., Min, J., & Yu, X. (2015). Full-color structured illumination optical sectioning microscopy. Scientific Reports, 5(1), 1–10. (e.g. “https://doi.org/10.1038/srep14513”)

Richter, A., Garcia, F. H., Keller, R. A., Billen, J., Economo, E. P., & Beutel, R. G. (2020). Comparative analysis of worker head anatomy of *Formica* and *Brachyponera* (Hymenoptera: Formicidae). Arthropod Systematics & Phylogeny, 78(1), 133–170. (e.g. “https://doi.org/10.26049/ASP78-1-2020-06”)

Nguyen, C. V., Lovell, D. R., Adcock, M., & La Salle, J. (2014). Capturing natural-colour 3D models of insects for species discovery and diagnostics. PloS one, 9(4), e94346. (e.g. “https://doi.org/10.1371/journal.pone.0094346”)

Sarnat, E. M., Garcia, F. H., Dudley, K., Liu, C., Fischer, G., & Economo, E. P. (2019). Ready species one: exploring the use of augmented reality to enhance systematic biology with a revision of Fijian *Strumigenys* (Hymenoptera: Formicidae). Insect Systematics and Diversity, 3(6), 6. (e.g. “https://doi.org/10.1093/isd/ixz005”)

Snodgrass, R. E. (1939). The Principles of Insect Physiology. Science, 90, 159–159. (e.g. “https://doi.org/10.1126/science.90.2329.159”)

Sosa, G. D., Rodríguez, S., Guaje, J., Victorino, J., Mejía, M., Fuentes, L. S., Ramirez, A., & Franco, H. (2016). 3D surface reconstruction of entomological specimens from uniform multi-view image datasets. [Paper presentation] XXI Symposium on Signal Processing, Images and Artificial Vision (STSIVA) (pp. 1-8). IEEE. (e. g. “https://doi.org/10.1109/STSIVA.2016.7743319”)

Ströbel, B., Schmelzle, S., Blüthgen, N., & Heethoff, M. (2018). An automated device for the digitization and 3D modelling of insects, combining extended-depth-of-field and all-side multi-view imaging. ZooKeys, 759, 1. (e.g. “https://doi.org/10.3897/zookeys.759.24584”)

Taylor, G. J., Hall, S. A., Gren, J. A., & Baird, E. (2020). Exploring the visual world of fossilized and modern fungus gnat eyes (Diptera: Keroplatidae) with X-ray microtomography. Journal of the Royal Society Interface, 17(163), 20190750. (e.g. “https://doi.org/10.1098/rsif.2019.0750”)

Unity Software, 2018.

van de Kamp, T., Schwermann, A. H., dos Santos Rolo, T., Lösel, P. D., Engler, T., Etter, W., … & Krogmann, L. (2018). Parasitoid biology preserved in mineralized fossils. Nature communications, 9(1), 1–14. (e.g. “https://doi.org/10.1038/s41467-018-05654-y”)

Yoder, M. J., Mikó, I., Seltmann, K. C., Bertone, M. A., & Deans, A. R. (2010). A gross anatomy ontology for Hymenoptera. PloS one, 5(12), e15991. (e.g. “https://doi.org/10.1371/journal.pone.0015991”)

Zilverschoon, M., Vincken, K. L., & Bleys, R. L. (2017). The virtual dissecting room: Creating highly detailed anatomy models for educational purposes. Journal of biomedical informatics, 65, 58–75. (e.g. “https://doi.org/10.1016/j.jbi.2016.11.005”)

Zimmermann, D., & Vilhelmsen, L. (2016). The sister group of Aculeata (Hymenoptera)–evidence from internal head anatomy, with emphasis on the tentorium. Arthropod Systematic Phylogeny, 74(2), 195–218.

