## Supplementary 01 for "EntomonVR: a New Virtual Reality Game for Learning Insect Morphology"

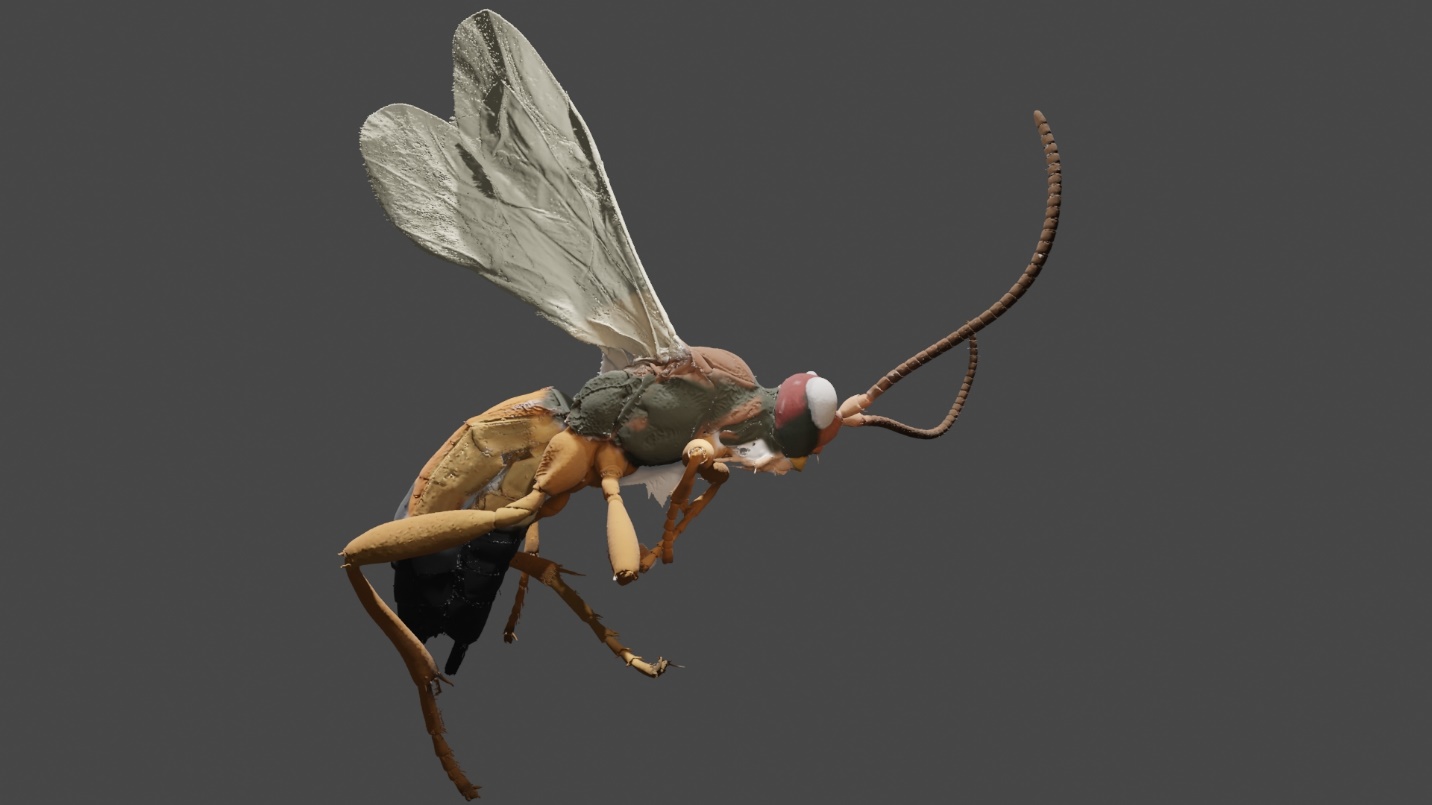

**Figure 1** 3D model of Aleiodes arnoldii Tobias, 1976 (Hymenoptera: Braconidae)

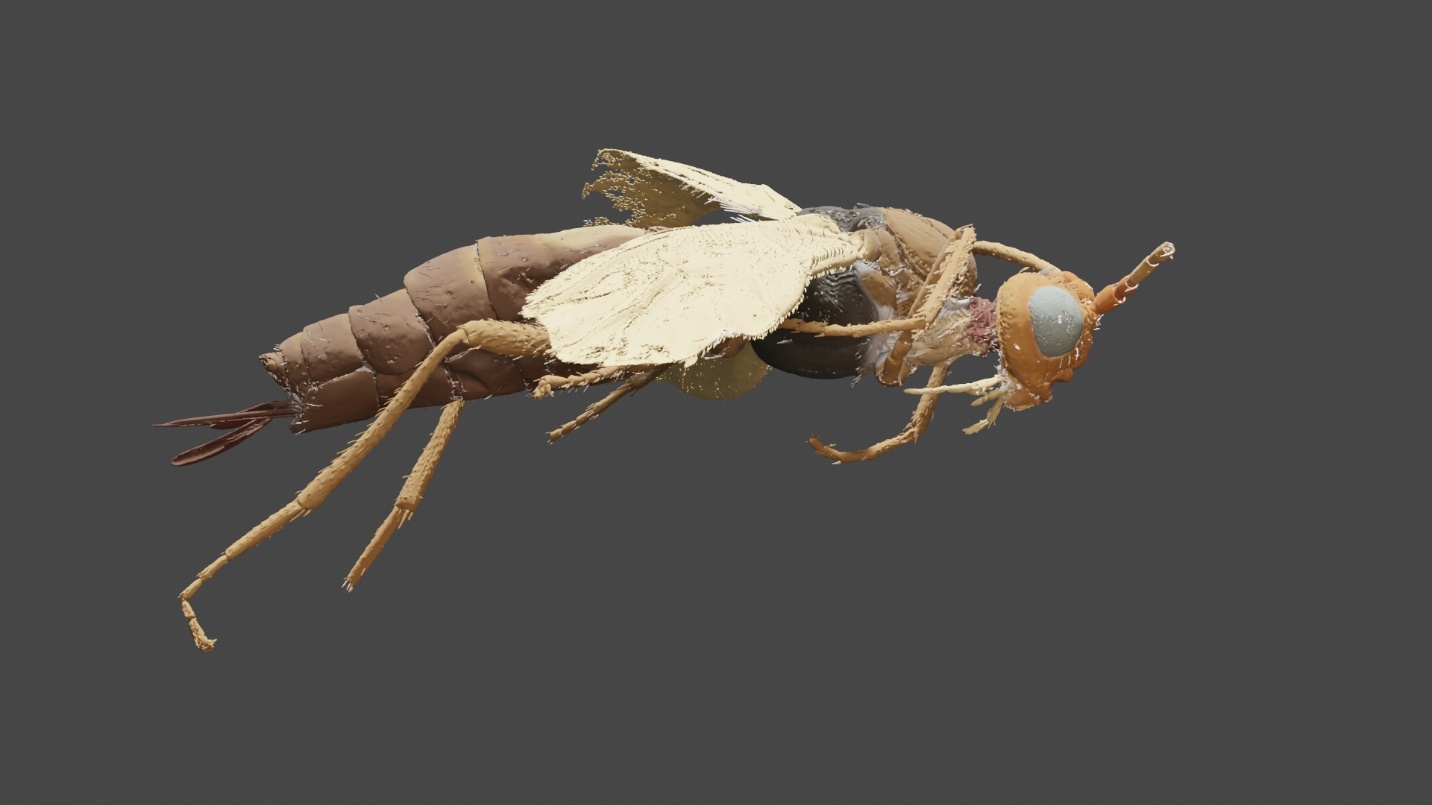

**Figure 2** 3D model of Hormius moniliatus Nees, 1811 (Hymenoptera: Braconidae)

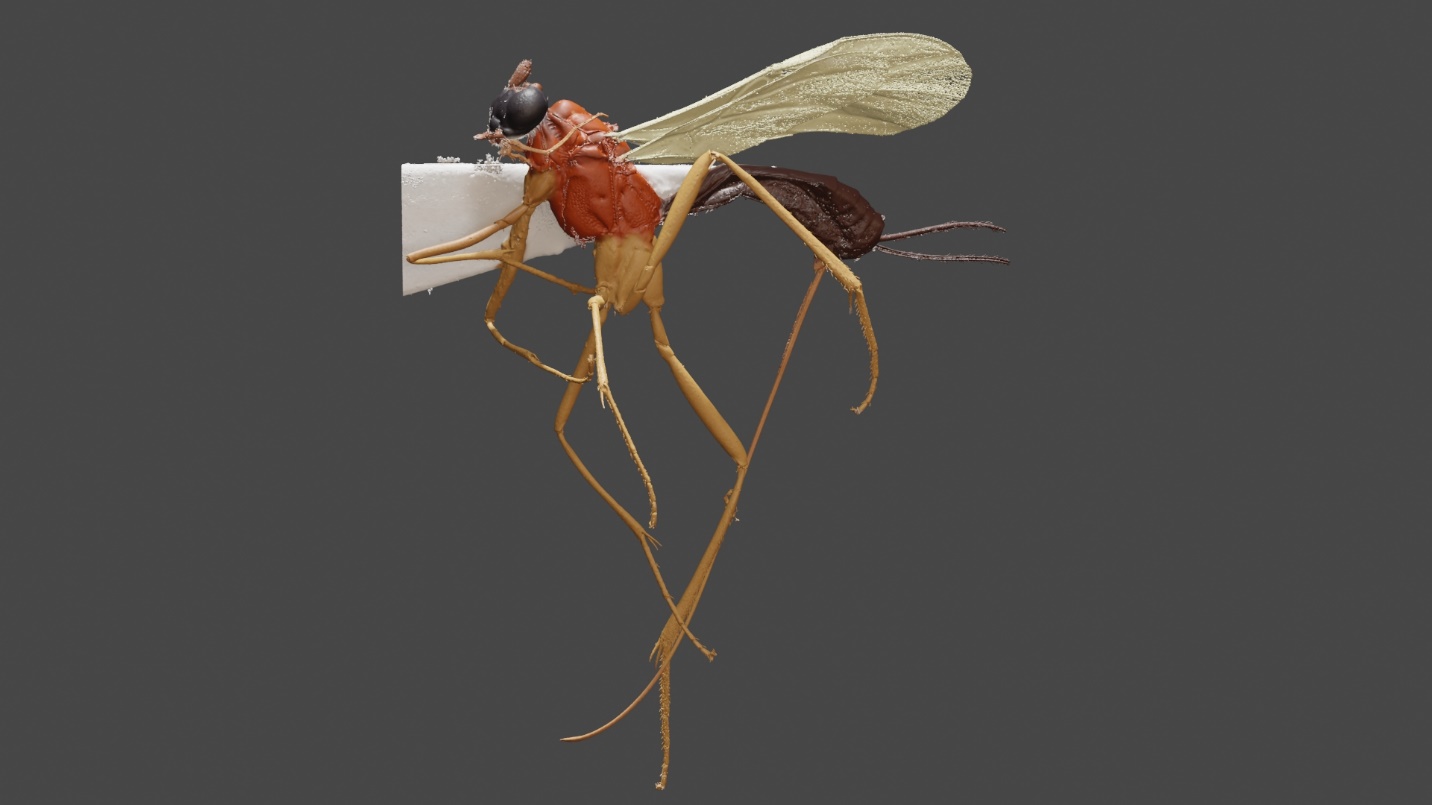

**Figure 3** 3D model of Macrocentrus bicolor Curtis, 1833 (Hymenoptera: Braconidae)

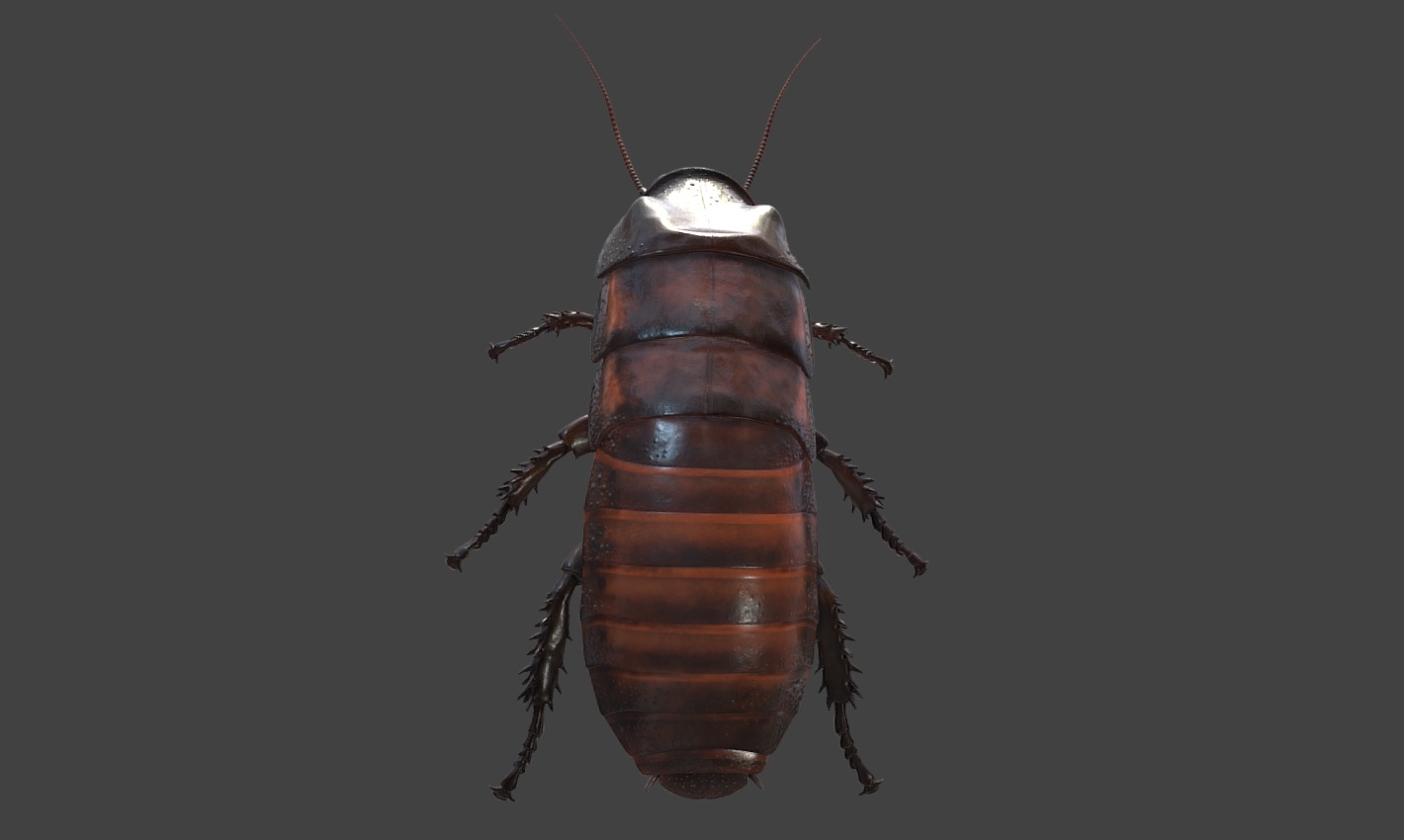

**Figure 4** 3D model of Gromphadorhina portentosa Schaum, 1853 (Blattodea: Blaberidae)

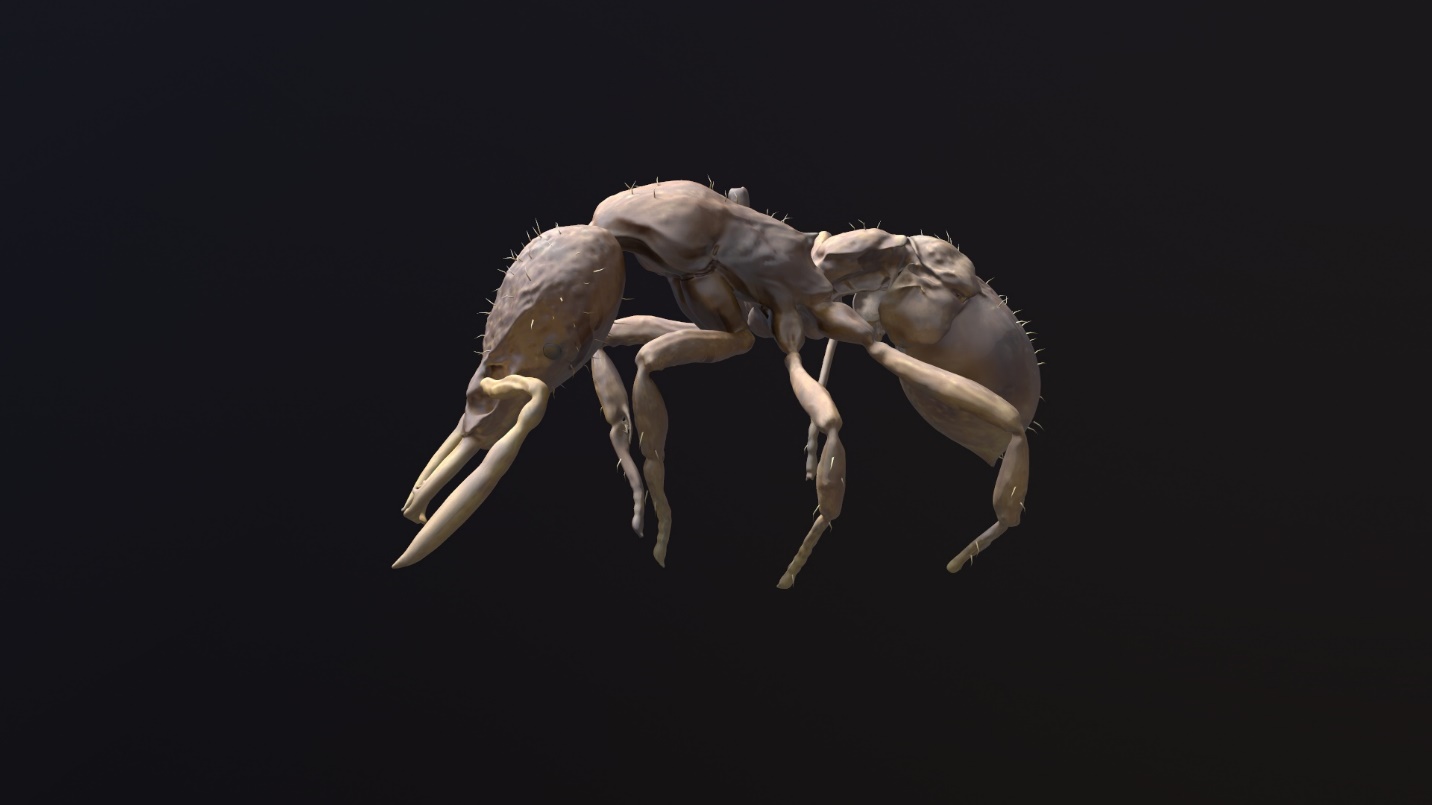

**Figure 5** 3D model of Strumigenys anorak Sarnat et al., 2019 (Hymenoptera: Formicidae)

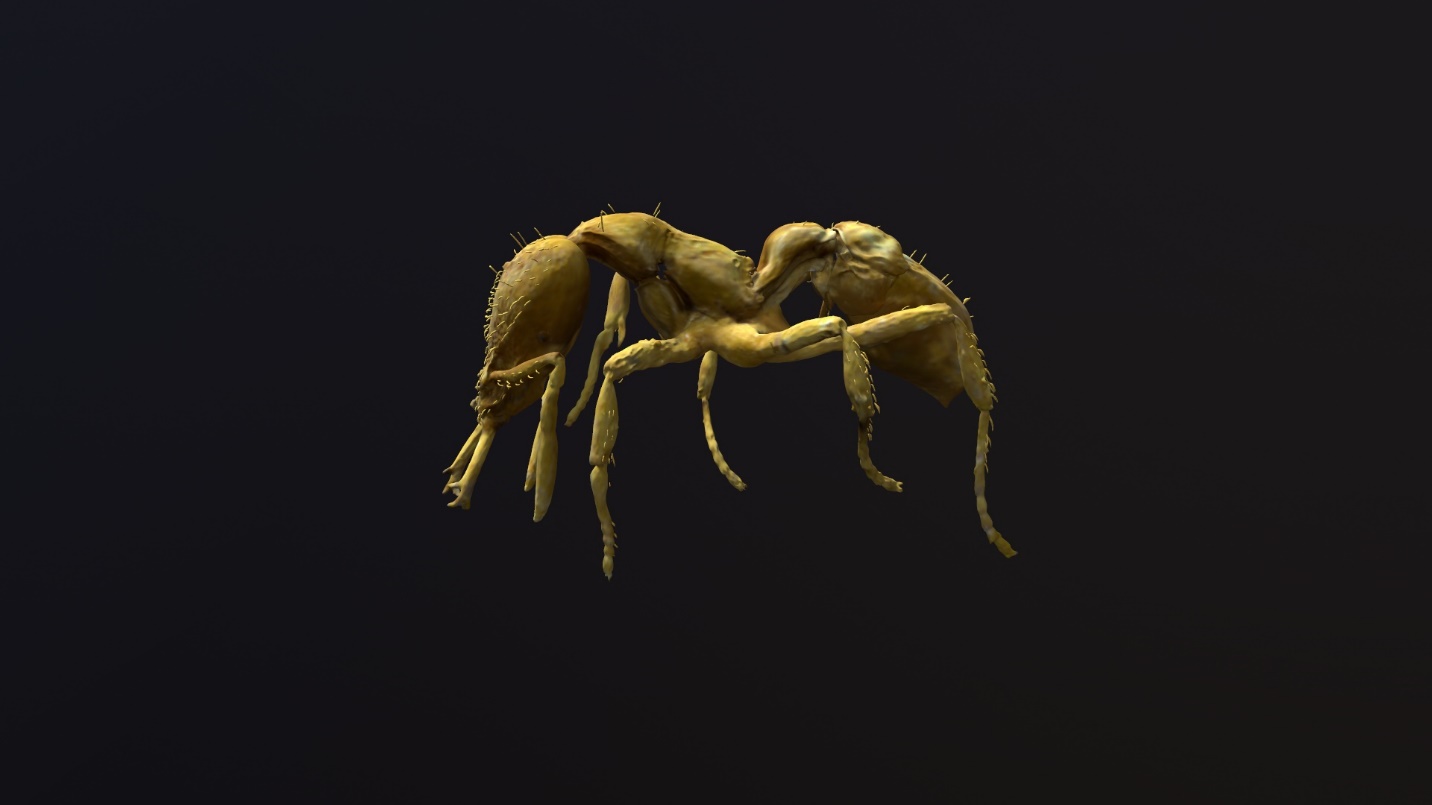

**Figure 6** 3D model of Strumigenys artemis Sarnat et al., 2019 (Hymenoptera: Formicidae)

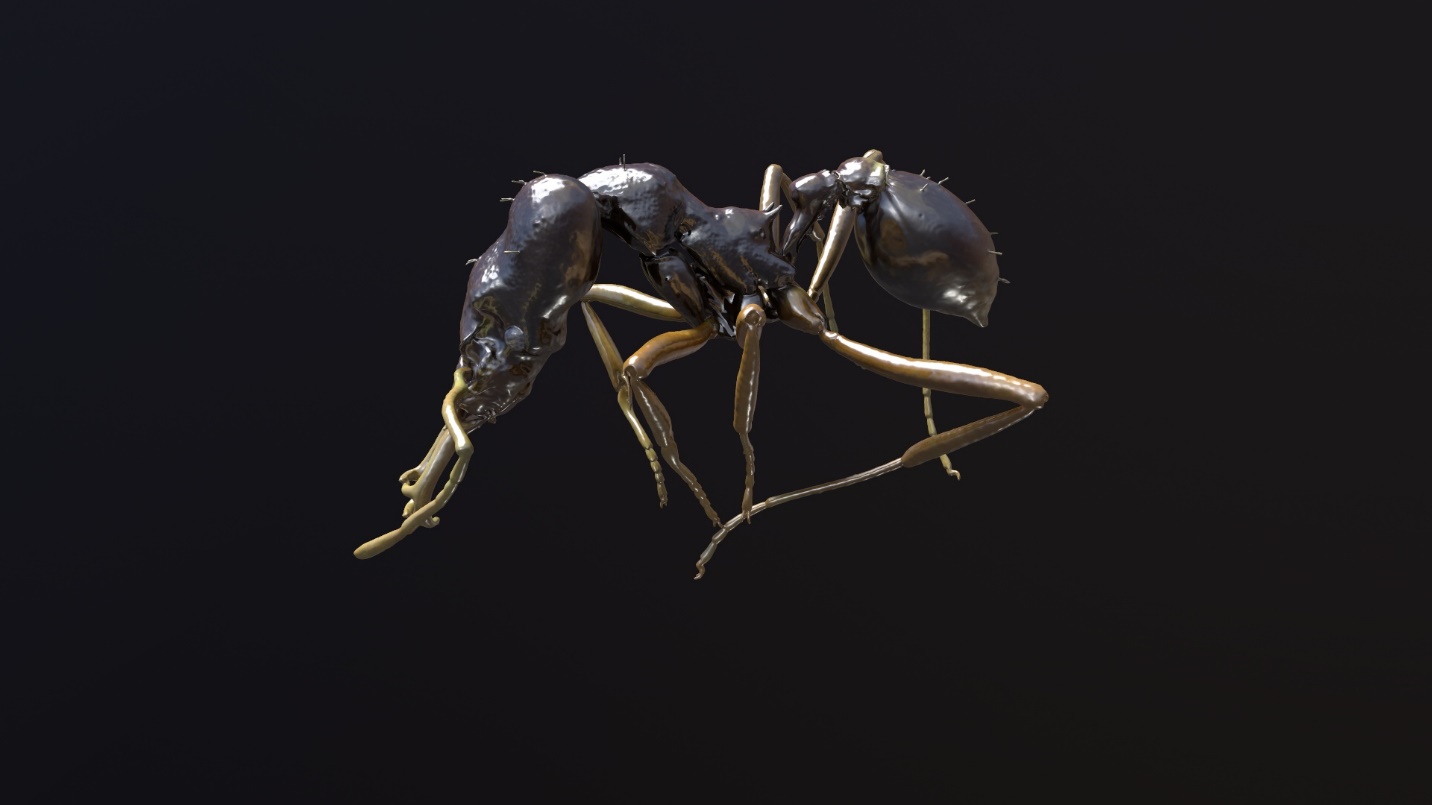

**Figure 7** 3D model of Strumigenys avatar Sarnat et al., 2019 (Hymenoptera: Formicidae)

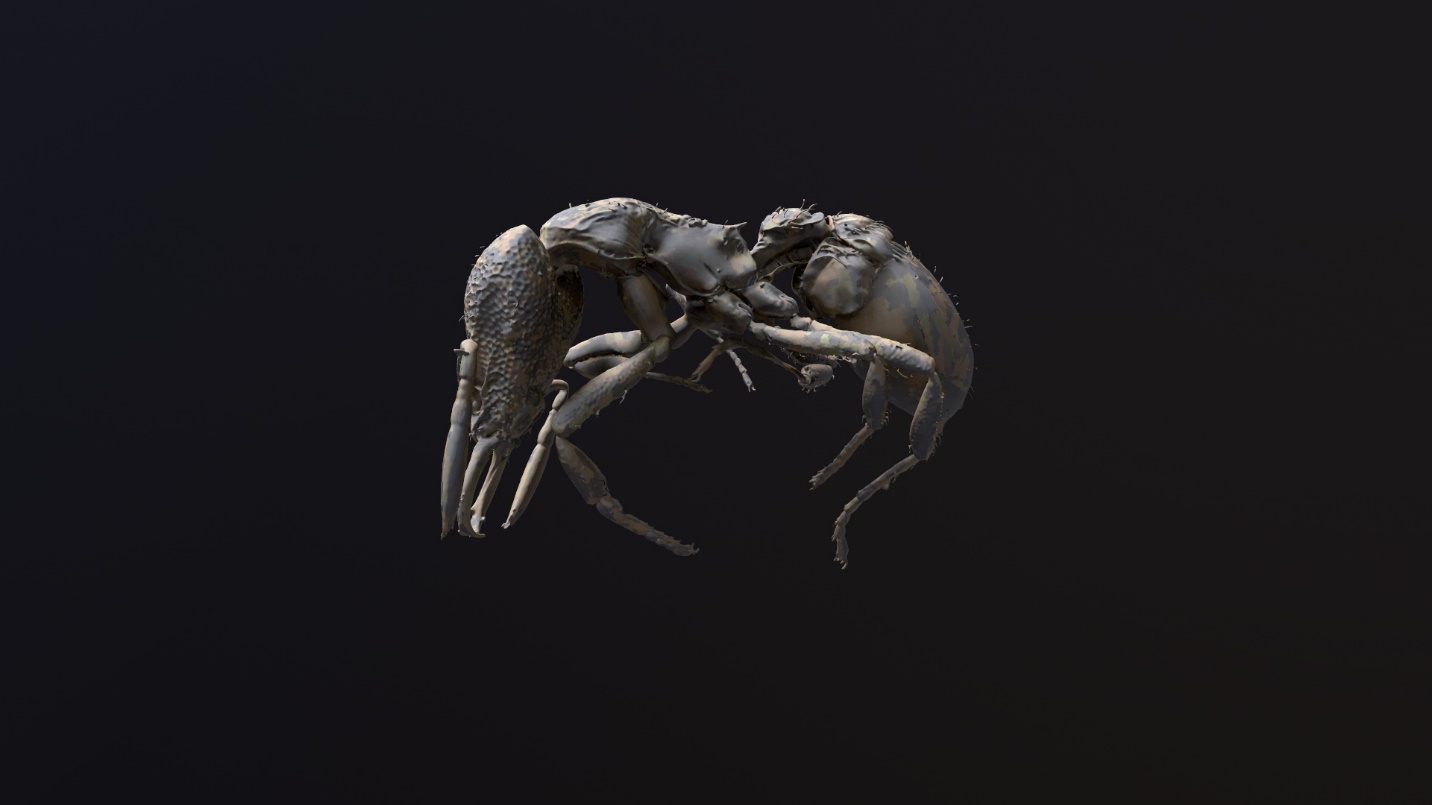

**Figure 8** 3D model of Strumigenys gunter Sarnat et al., 2019 (Hymenoptera: Formicidae)

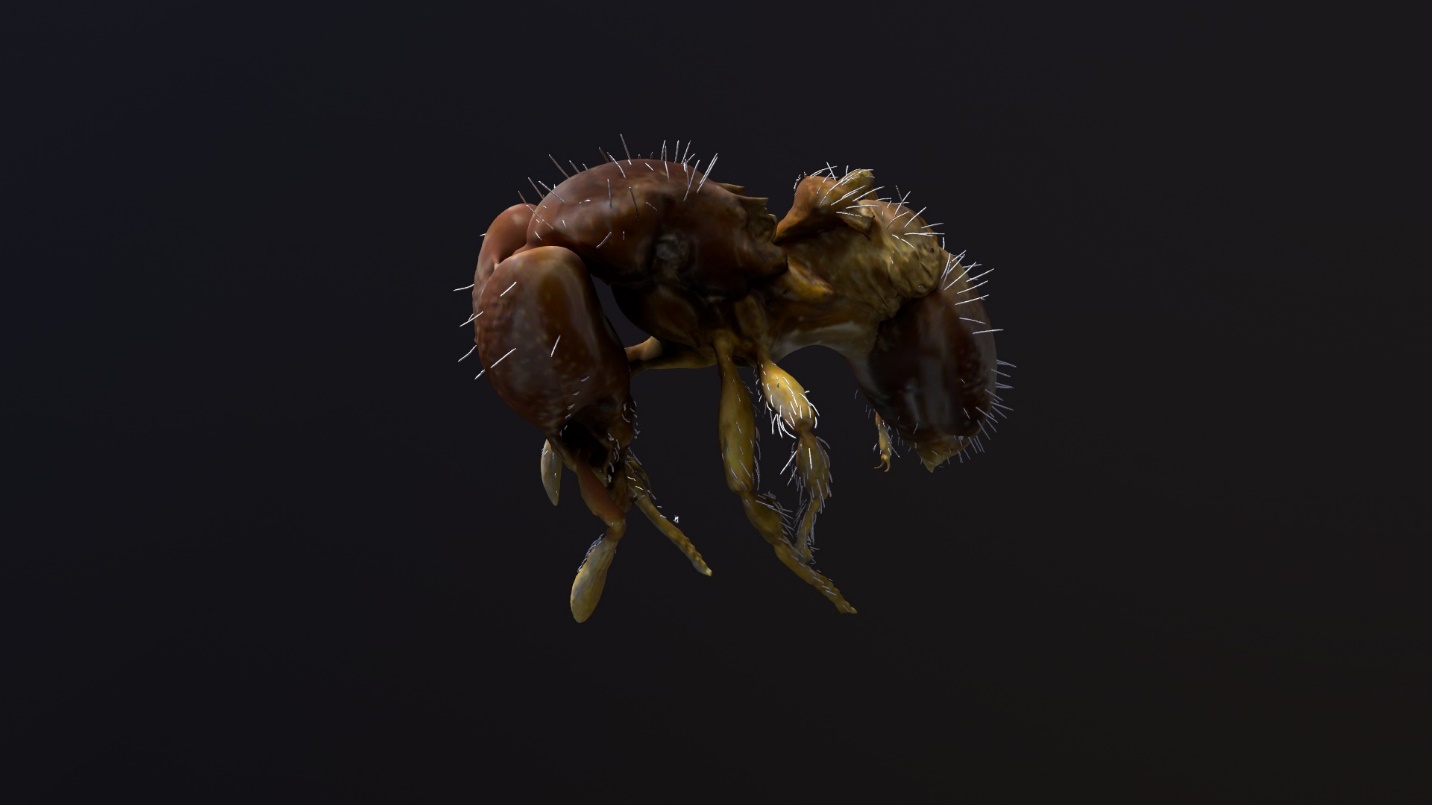

**Figure 9** 3D model of Strumigenys oasis Sarnat et al., 2019 (Hymenoptera: Formicidae)

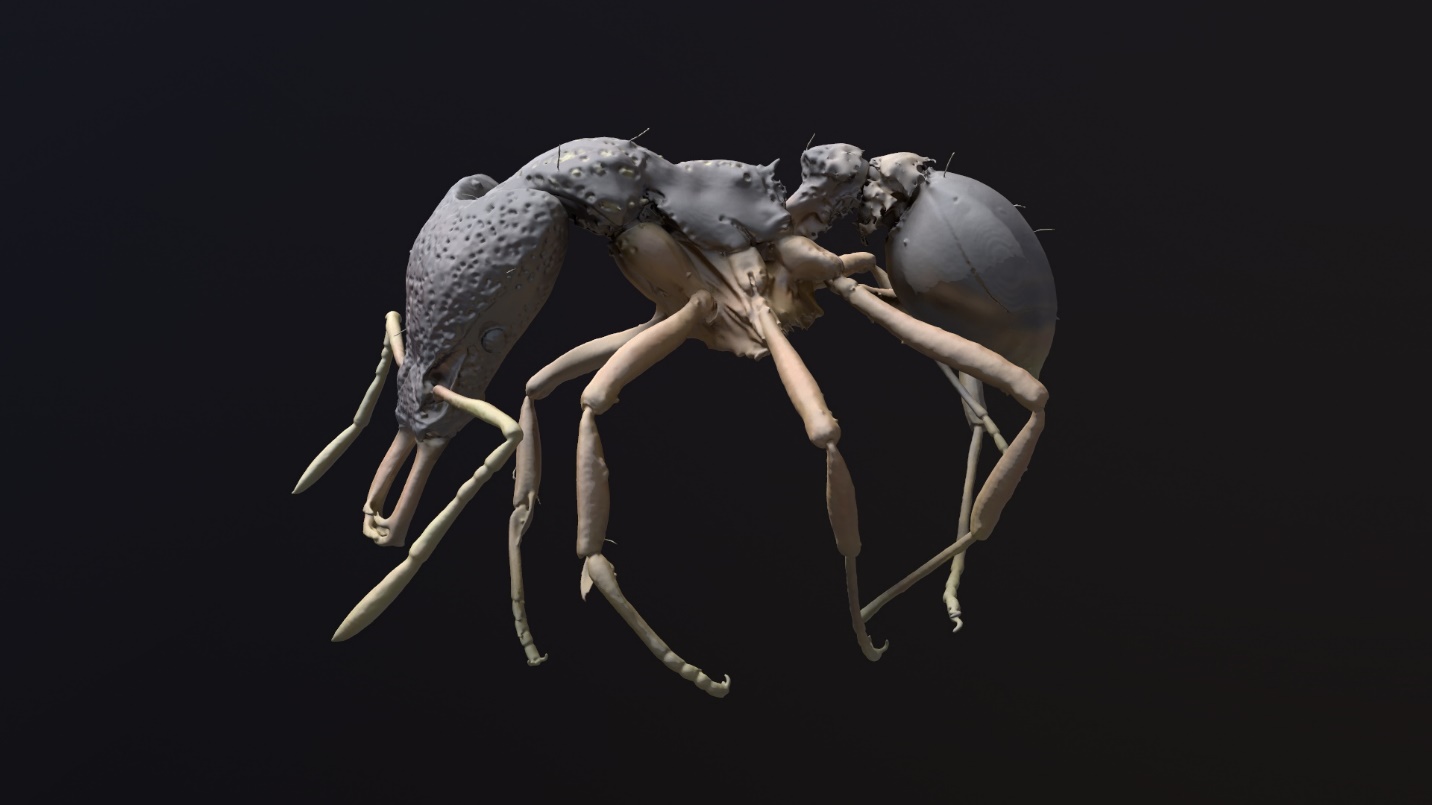

**Figure 10** 3D model of Strumigenys parzival Sarnat et al., 2019 (Hymenoptera: Formicidae)

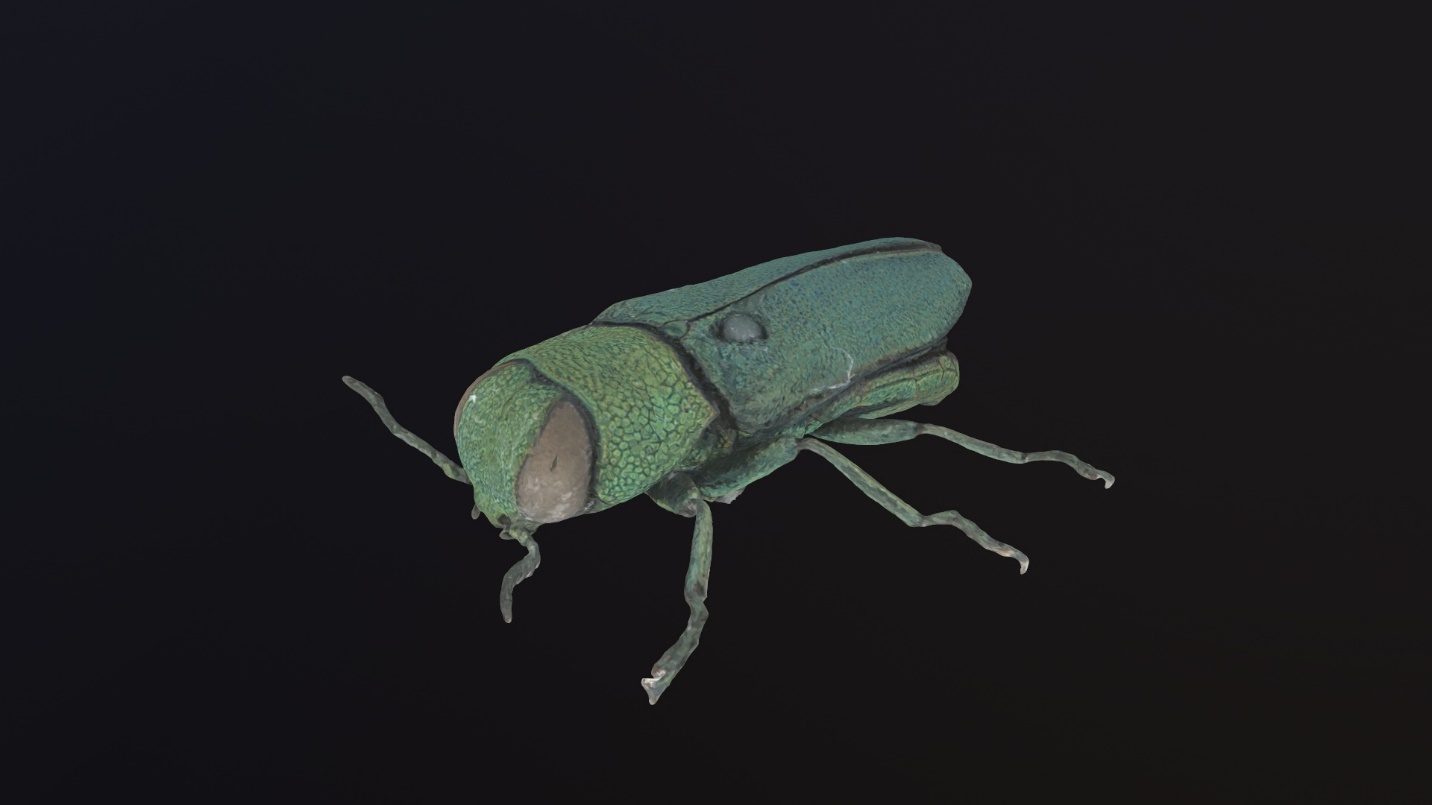

**Figure 11** 3D model of Anthaxia nitidula Linnaeus, 1758 (Coleoptera: Buprestidae)

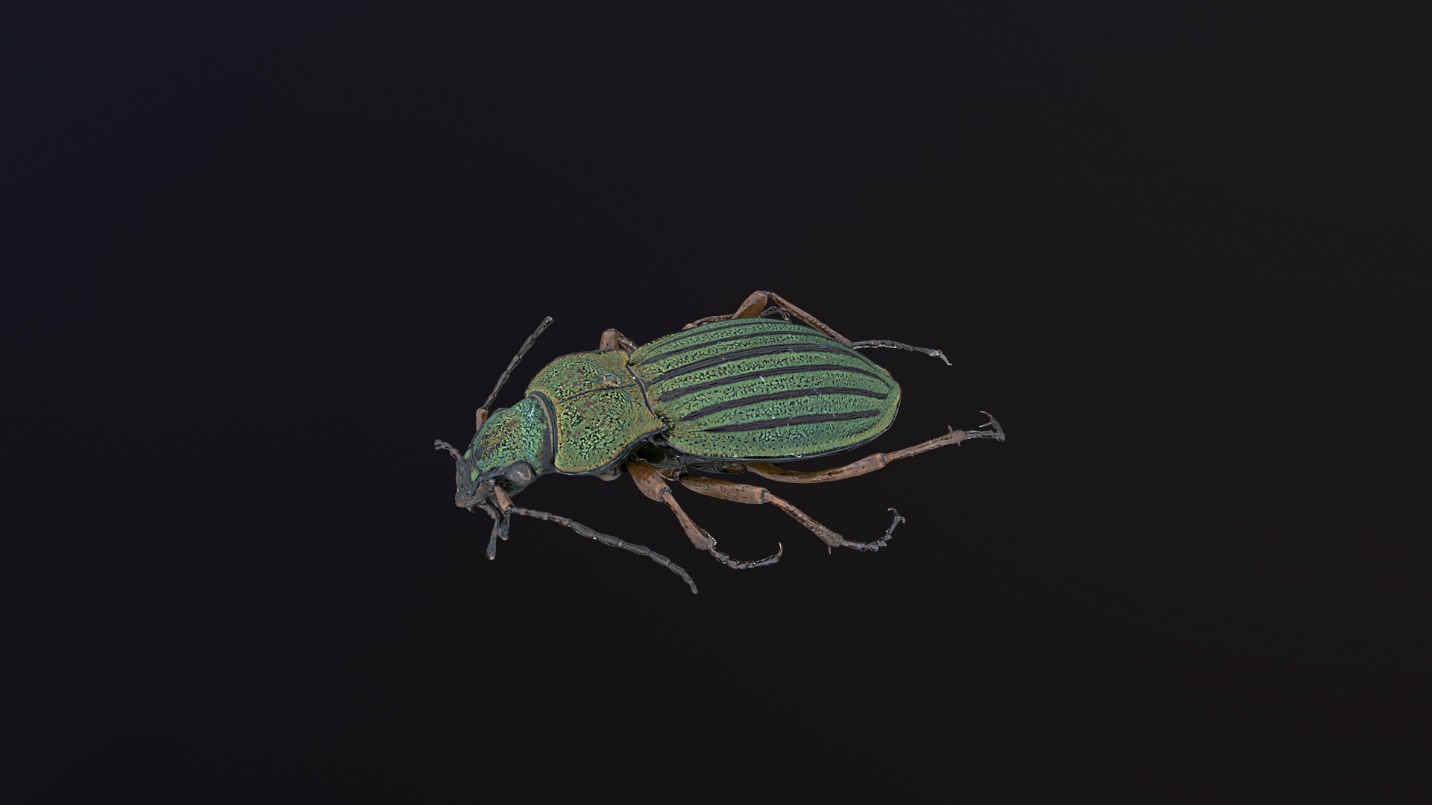

**Figure 12** 3D model of Carabus auronitens Fabricius, 1792 (Coleoptera: Carabidae)

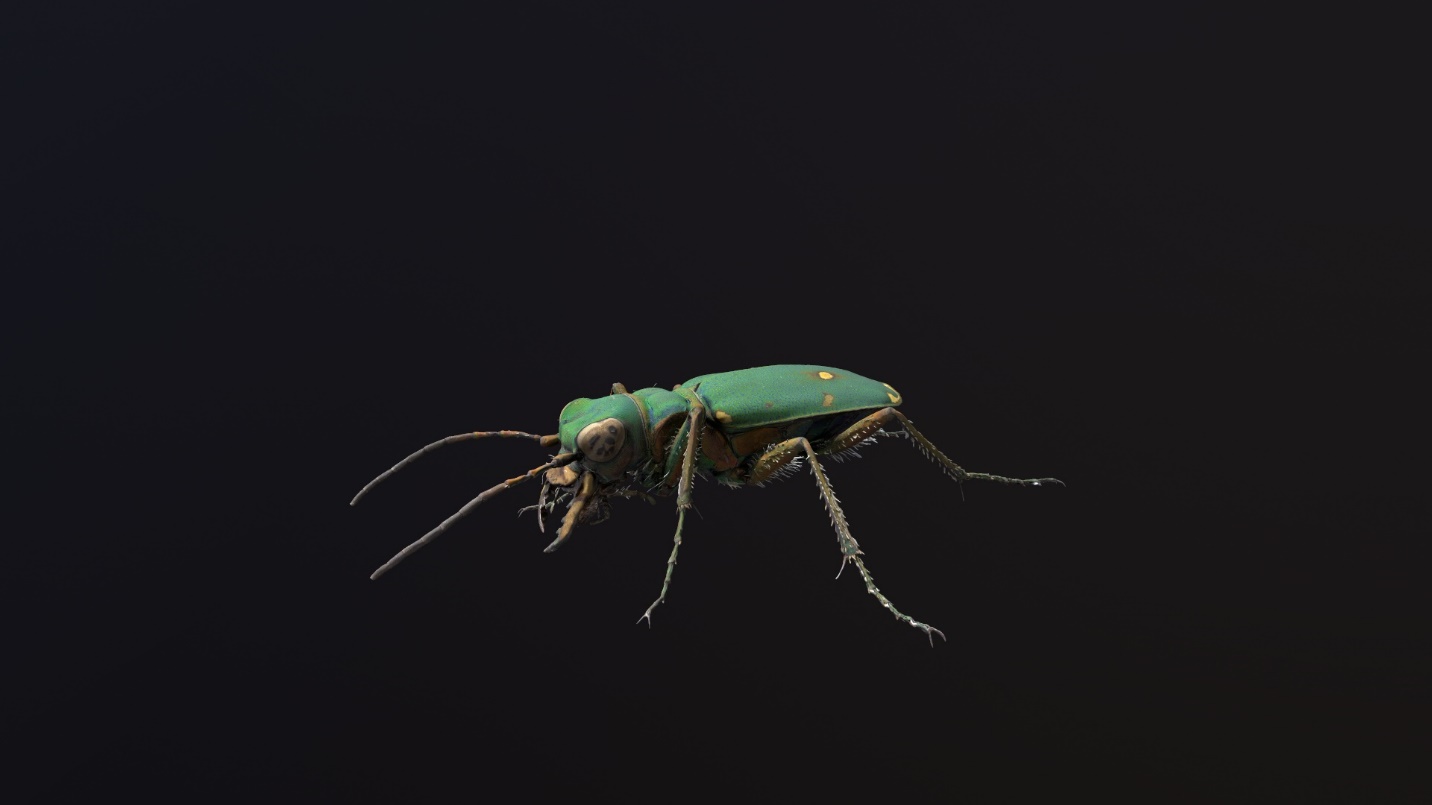

**Figure 13** 3D model of Cicindela campestris Linnaeus, 1758 (Coleoptera: Carabidae)

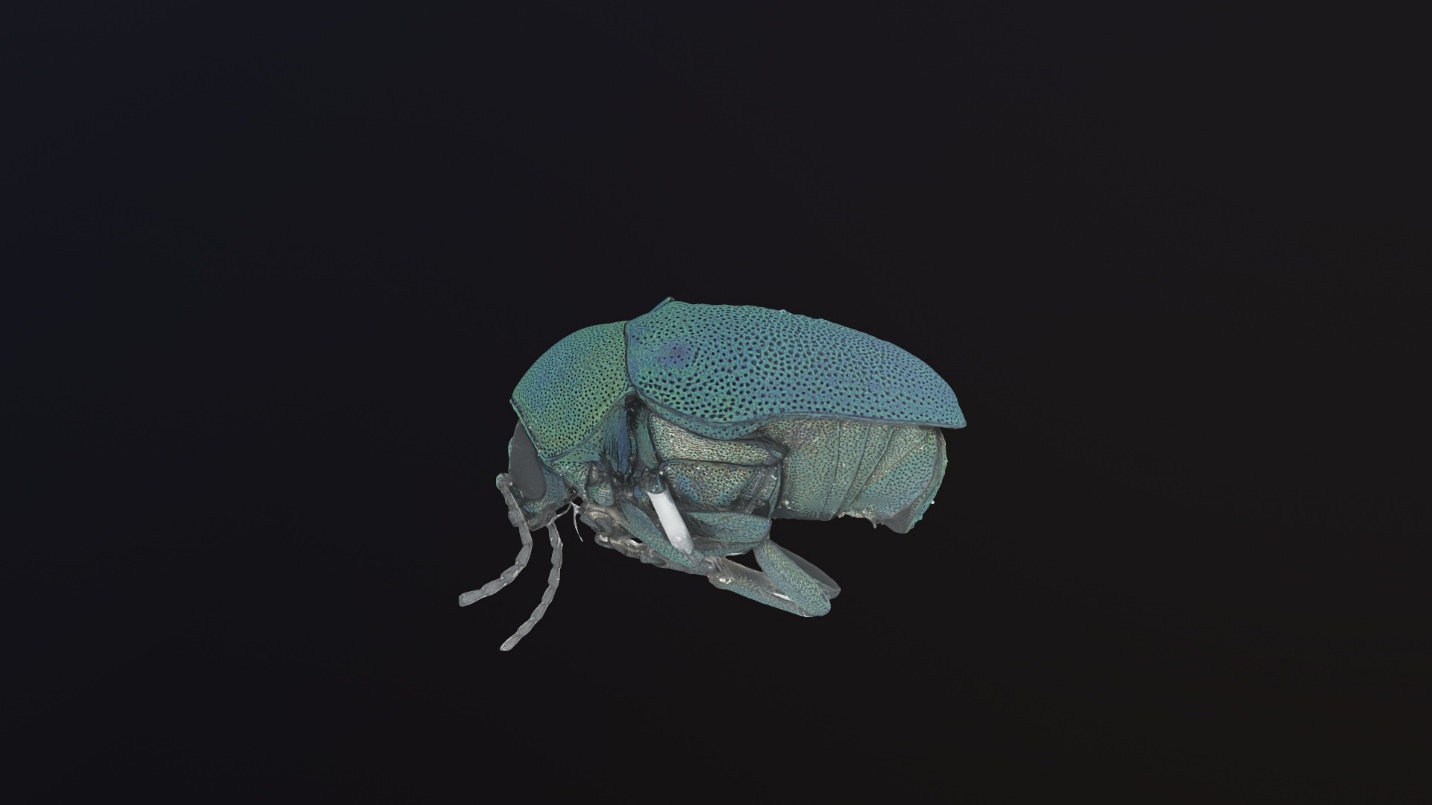

**Figure 14** 3D model of Cryptocephalus sericeus Linnaeus, 1758 (Coleoptera: Chrysomelidae)

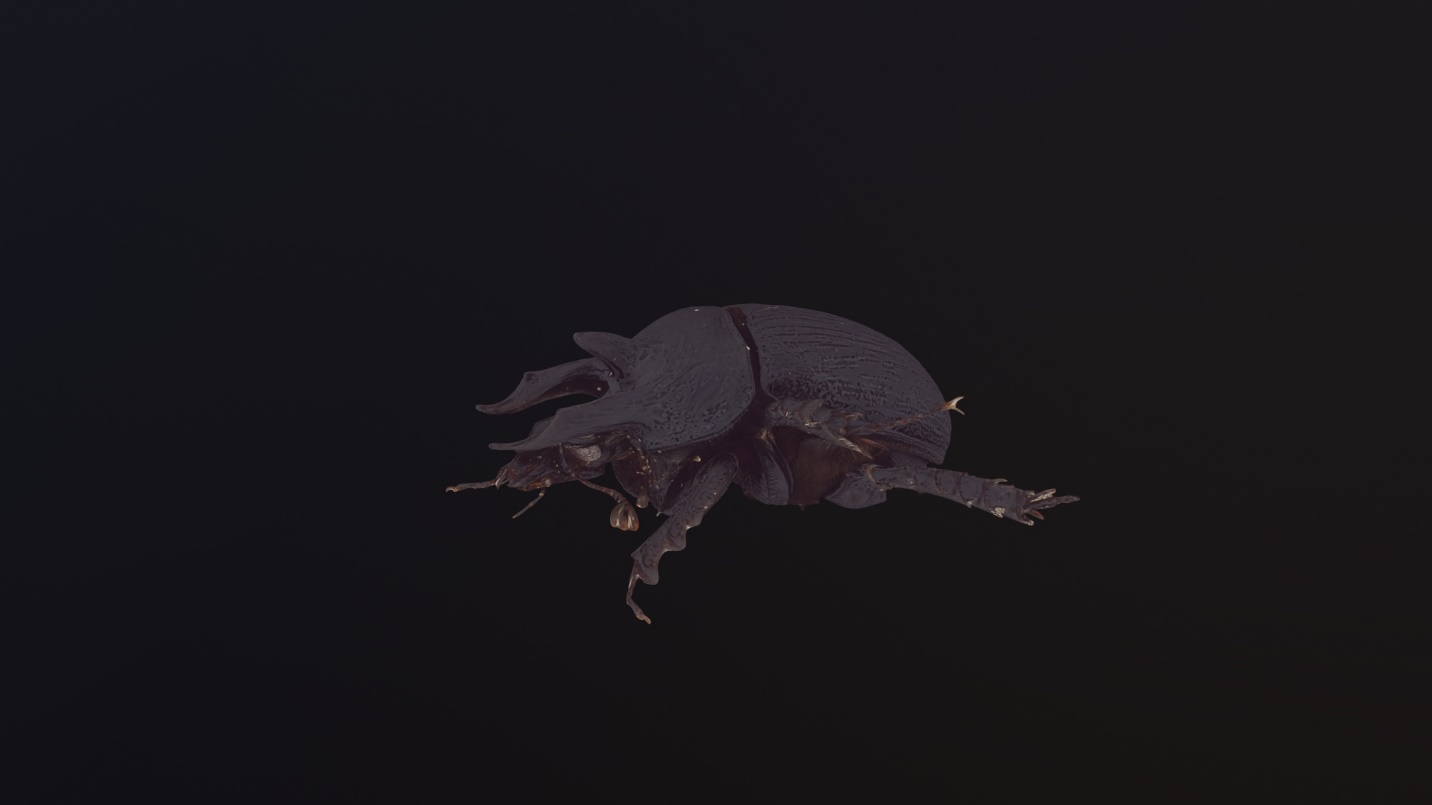

**Figure 15** 3D model of Typhaeus typhoeus Linnaeus, 1758 (Coleoptera: Geotrupidae)

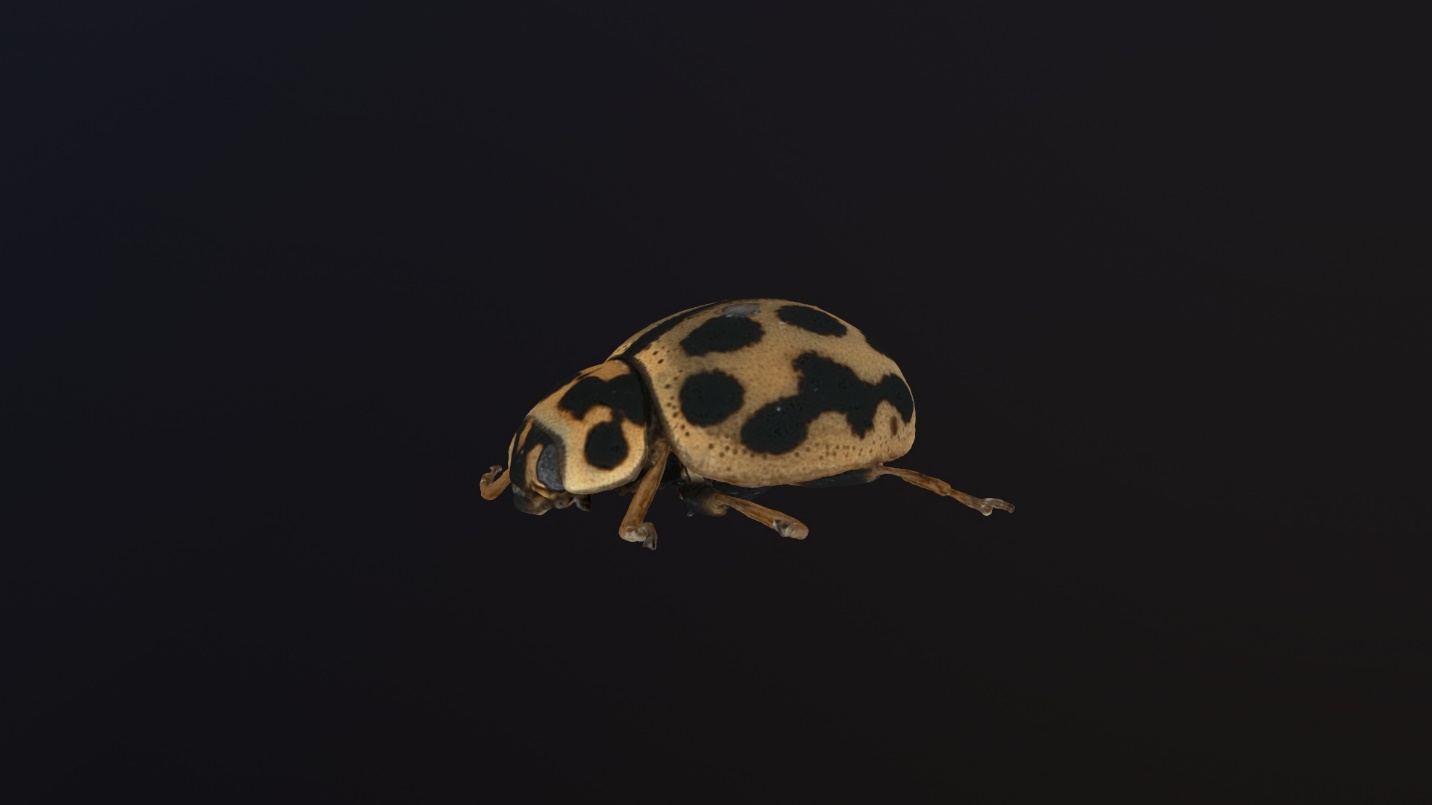

**Figure 16** 3D model of Tytthaspis sedecimpunctata Linnaeus, 1761 (Coleoptera: Coccinellidae)

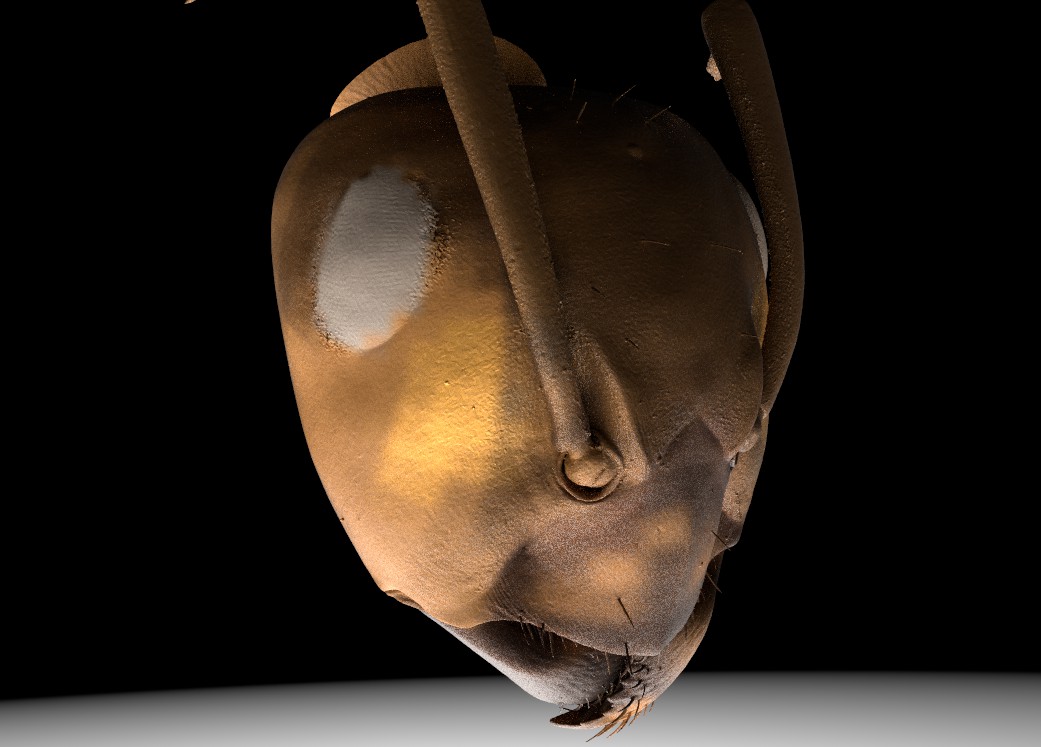

Figure 17 3D head model of Formica sp. (Hymenoptera: Formicidae)

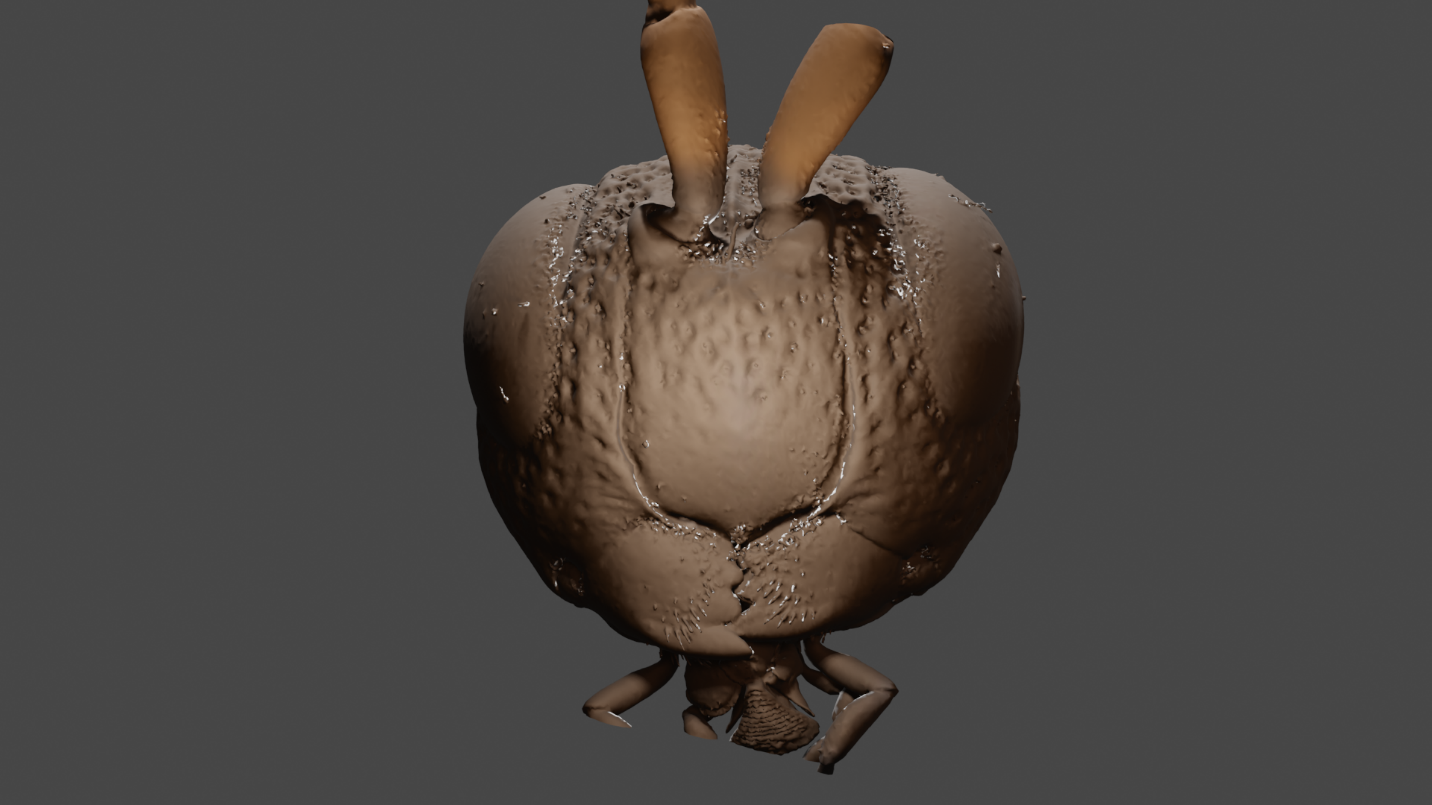

**Figure 18** 3D head model of Evaniella semaeoda Bradley, 1908 (Hymenoptera: Evaniidae)

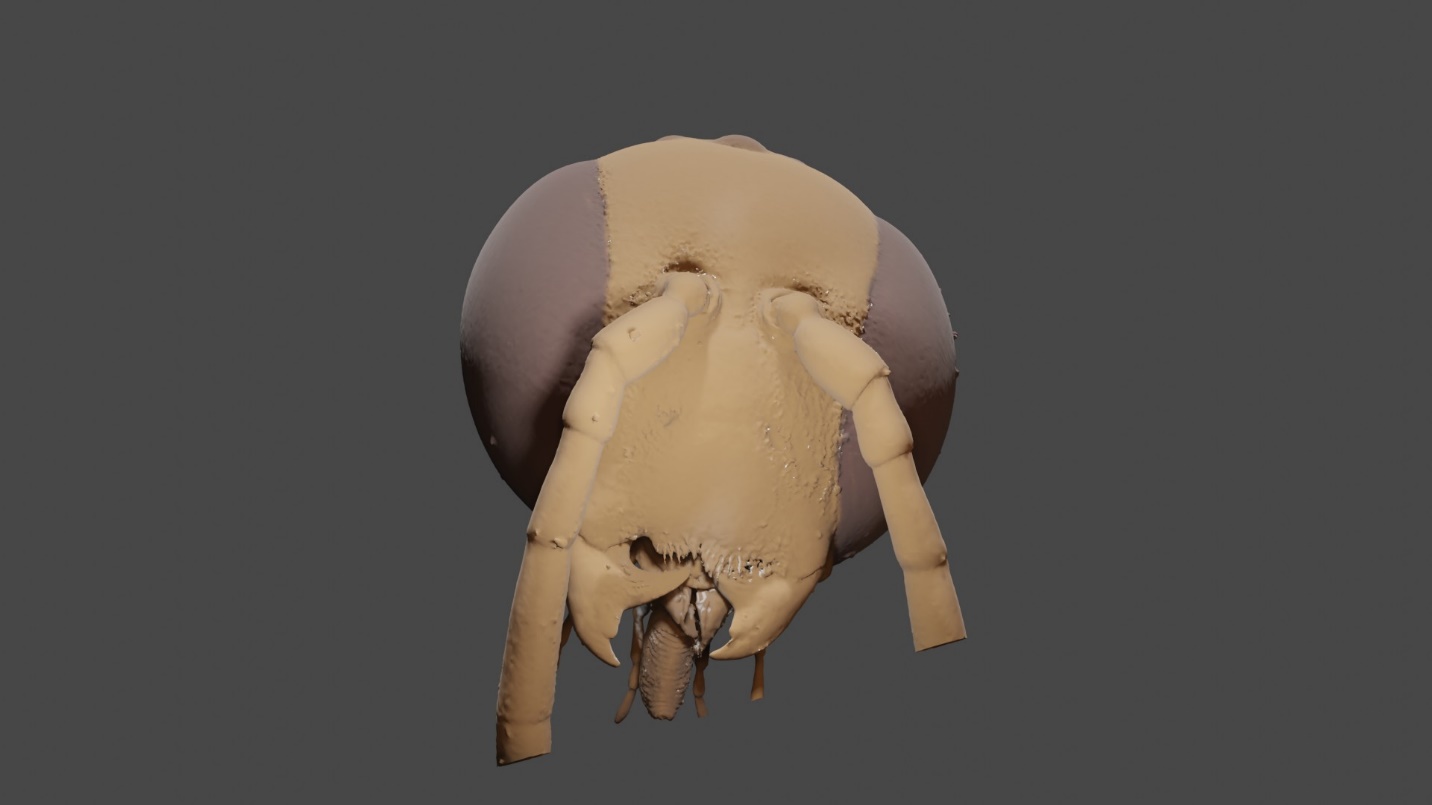

**Figure 19** 3D head model of Gasteruption tarsatorius Say, 1824 (Hymenoptera: Gasteruptiidae)

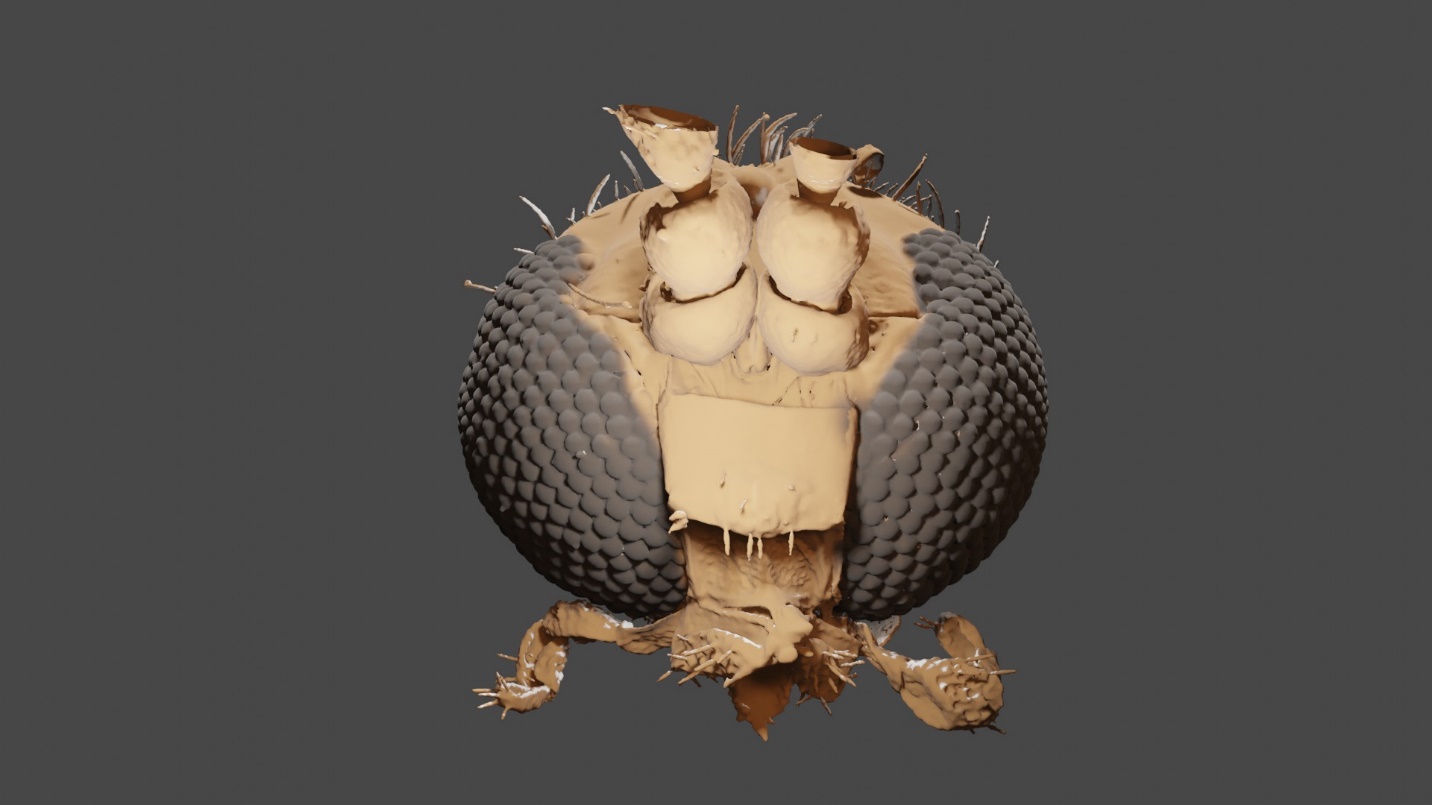

**Figure 20** 3D head model of Neoplatyura modesta Winnertz, 1863 (Diptera: Keroplatidae)

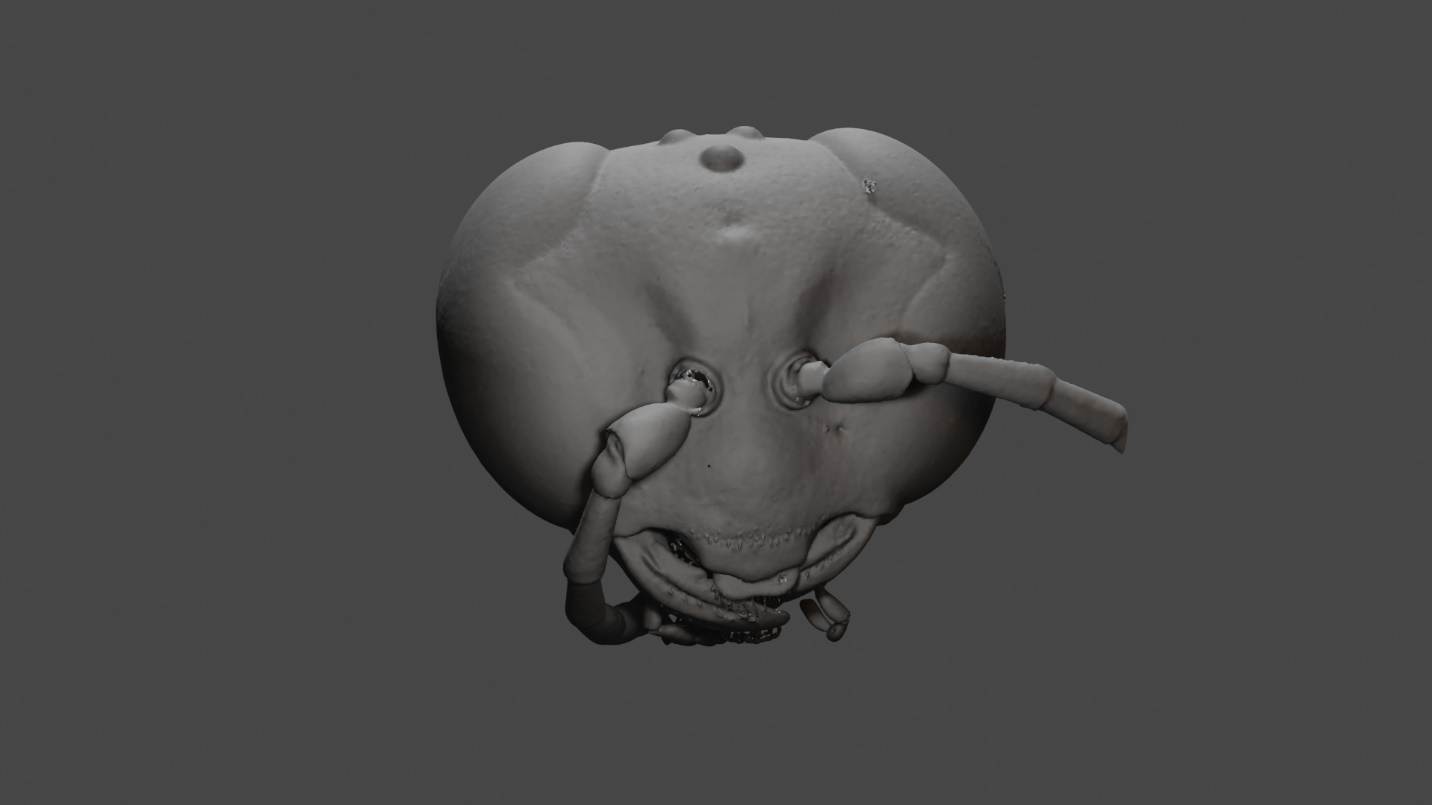

**Figure 21** 3D head model of Pison chilense Spinola, 1851 (Hymenoptera: Crabronidae)

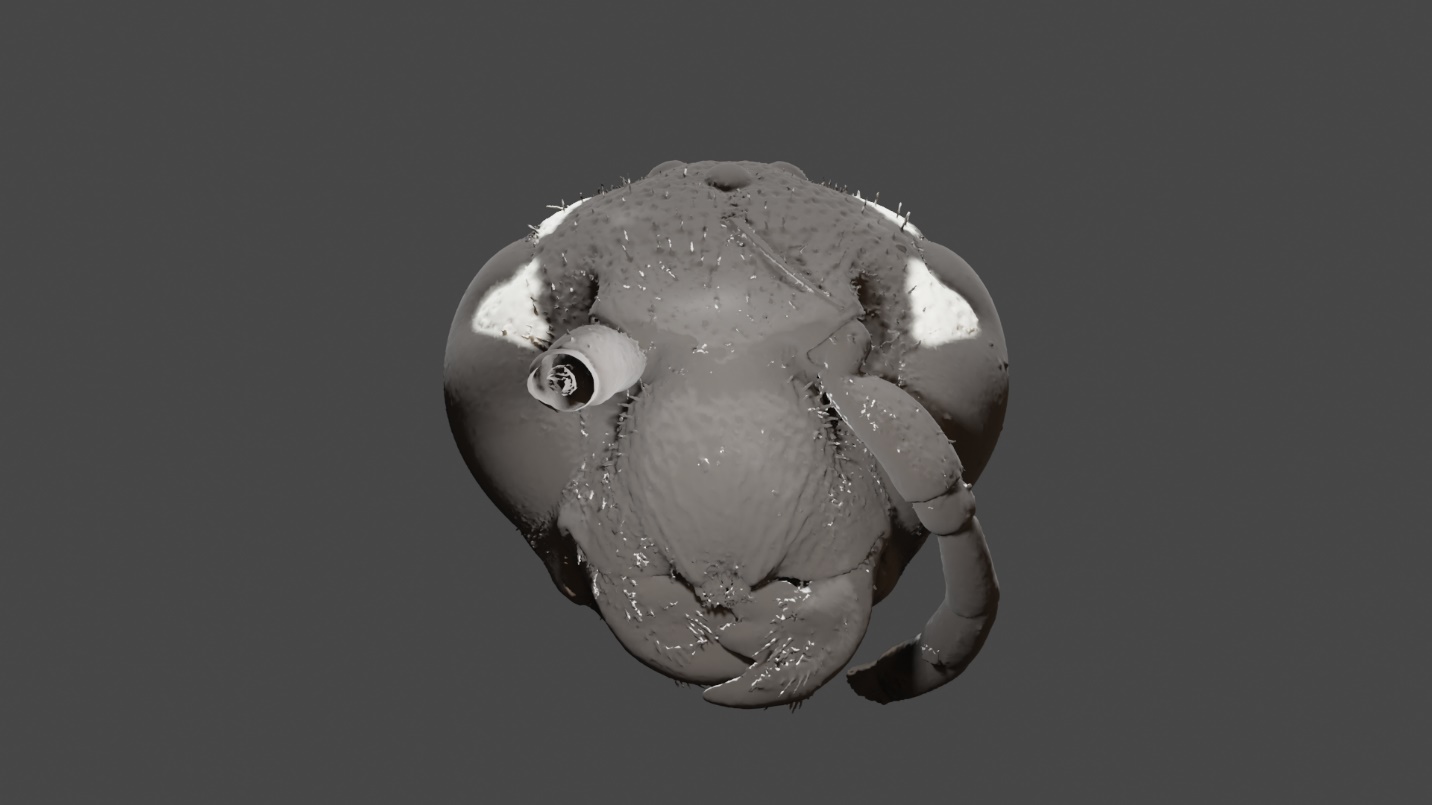

**Figure 22** 3D head model of Sapyga pumila Cresson, 1880 (Hymenoptera: Sapygidae)

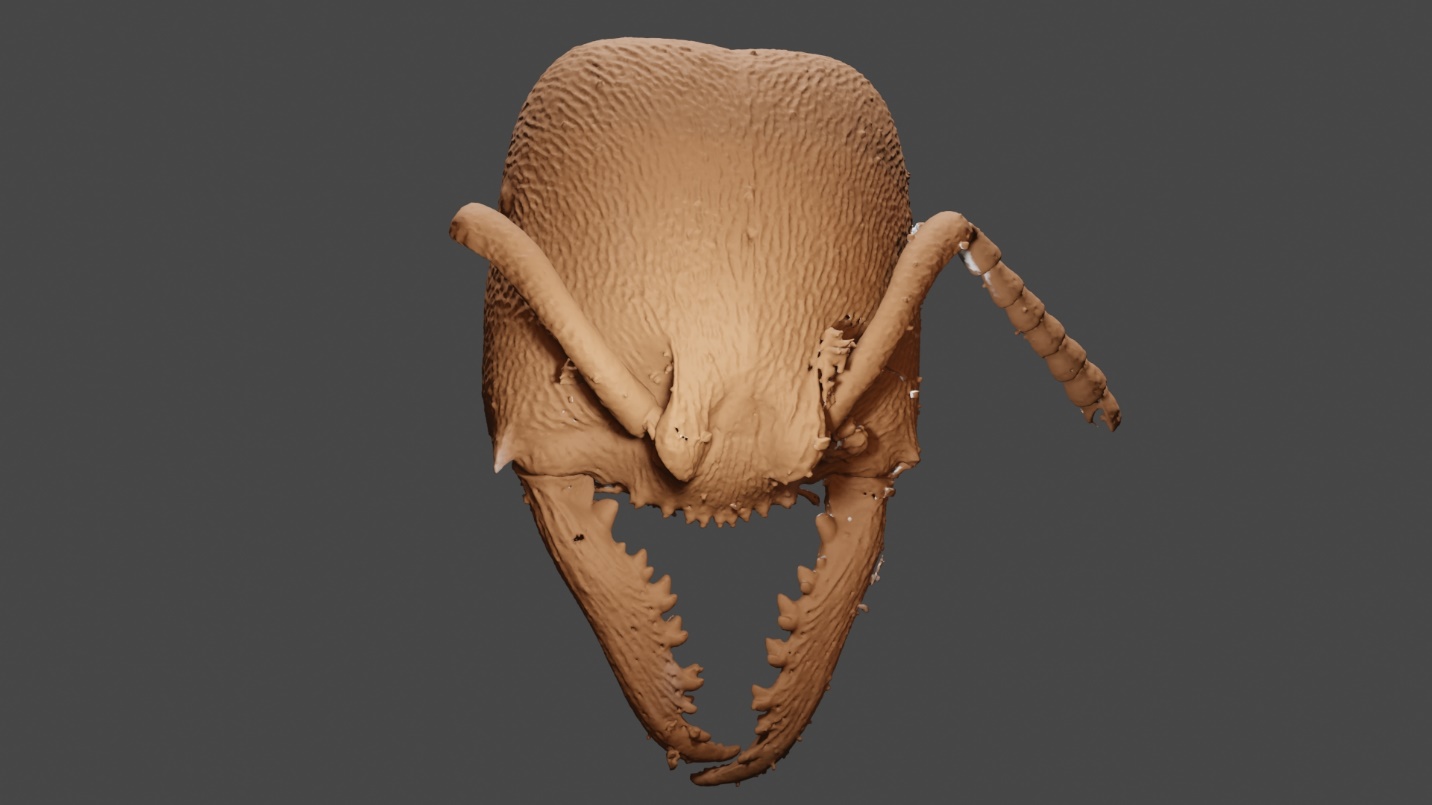

**Figure 23** 3D head model of Stigmatomma pallipes Haldeman, 1844 (Hymenoptera: Formicidae)

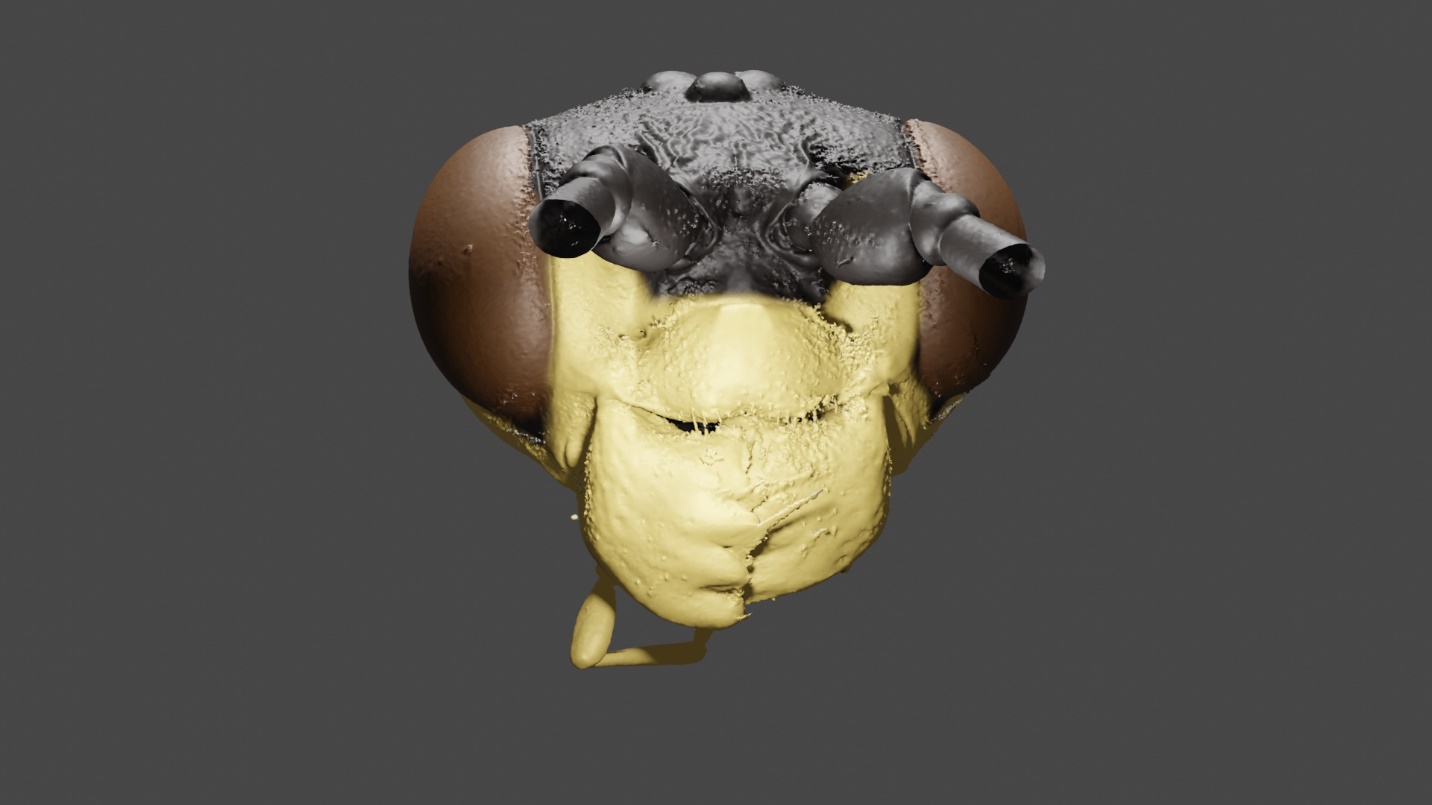

**Figure 24** 3D head model of Orthogonalys pulchella Cresson, 1867 (Hymenoptera: Trigonalyidae)

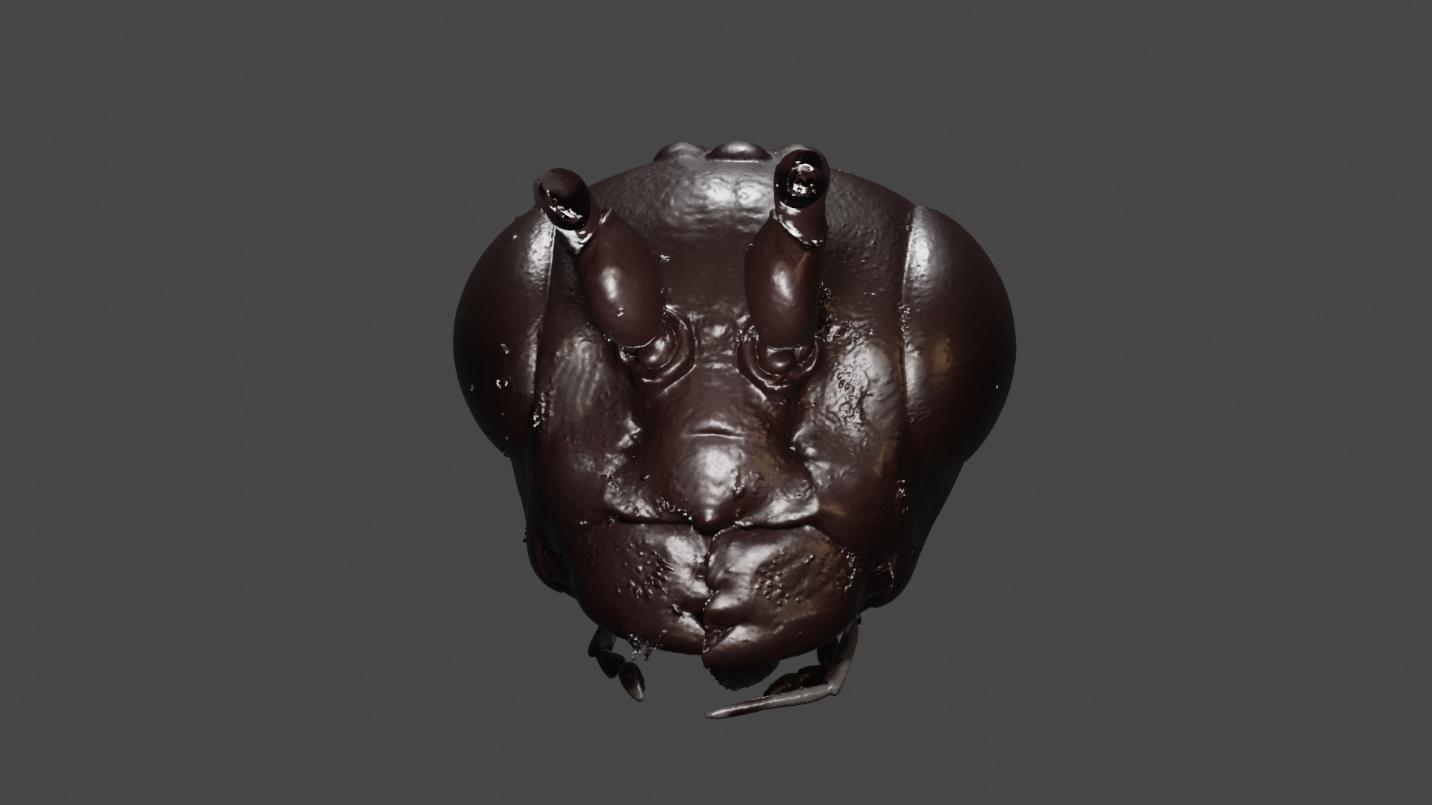

**Figure 25** 3D head model of Pristaulacus strangaliae Rohwer, 1917 (Hymenoptera: Aulacidae)
